# Convergent Cellular Adaptation to Freeze-Thaw Stress via a Quiescence-like State in Yeast

**DOI:** 10.1101/2024.11.26.625389

**Authors:** Charuhansini Tvishamayi, Farhan Ali, Nandita Chaturvedi, Nithila Madhu-Kumar, Zeenat Rashida, Chandan Muni Reddy, Ankita Ray, Stephan Herminghaus, Shashi Thutupalli

**Author notes:** Equal Contribution, listed alphabetically (Farhan Ali, Nandita Chaturvedi).

## Abstract

Extreme stress, such as freeze-thaw, poses a severe challenge to many organisms, but the mechanisms underlying their adaptation to survive such stress remain elusive. Here, we show that *Saccharomyces cerevisiae* can rapidly evolve freeze-thaw tolerance through a physiological state transition, with survival increasing nearly two orders of magnitude from ≈2% to ≈70% in about 25 cycles of stress exposure. Evolved yeast cells exhibit a quiescence-like state, characterized by altered cellular physiology: increased intracellular trehalose accumulation, reduced membrane damage, cytoplasmic stiffening and an exit from a proliferative cycle. This mechano-chemically reinforced survival strategy emerges across independent evolutionary lines despite distinct genetic backgrounds, suggesting a convergent mechanism of adaptation. By integrating experimental evolution, biophysical measurements, genomic analysis, and a quantitative model that captures the adaptation dynamics, we reveal that stress tolerance can arise via a potentially generalizable, physiologically mediated adaptation strategy. These findings provide new insights into microbial survival under extreme conditions and suggest broader implications for cellular stress responses beyond yeast.

## INTRODUCTION

Cells across many domains of life survive extreme environmental perturbations, including complete desiccation and freezing, which can cause severe cellular damage. Cellular damage occurs via many mechanisms such as protein denaturation [1, 2], membrane rupture [3, 4], and oxidative stress [5]. To survive such conditions, organisms have evolved diverse adaptive mechanisms [6, 7]. Classical evolutionary models primarily emphasize genetic mutations as the basis for the evolution of such adaptations [8–10]. However, recent evidence suggests that cells can also adapt via phenotypic plasticity and physiological state transitions, allowing rapid, non-genetic survival advantages [11–16].

One widely recognized mechanism of stress resistance is the accumulation of protective molecules such as trehalose, which stabilizes proteins and membranes during dehydration and freezing [14, 17, 18]. Additionally, molecules such as trehalose have also been associated with transition to quiescence, a reversible state of metabolic suppression, to endure unfavorable conditions, in many microbes and multicellular organisms. Among many other characteristics, quiescence is characterized by increased cytoplasmic viscosity, enhanced resistance to oxidative damage, and improved long-term viability [19–22]. In microbes, quiescence has been linked to persistence under starvation and antimicrobial stress, while in multicellular organisms, it plays a role in stem cell maintenance and tumor dormancy [23–25].

Here, we investigate whether repeated exposure to extreme freeze-thaw stress selects for a quiescence-like adapted state in *Saccharomyces cerevisiae*. Using experimental evolution, we subjected yeast populations to repeated cycles of rapid freezing and thawing, a severe mechano-chemical perturbation that initially results in very low survival (*<*2%) [26, 27]. Strikingly, within ∼25 selection cycles, survival increased nearly 40-fold. The adapted yeast cells exhibit an increased prevalence of a quiescence-like state, characterized by altered cellular physiology: increased intracellular trehalose accumulation, reduced membrane damage, cytoplasmic stiffening and an exit from a proliferative cycle. Whole-genome sequencing revealed that the adaptation was not driven by unique genetic pathways but rather by multiple, distinct genetic backgrounds, all converging on the same mechano-chemically reinforced survival phenotype. Our findings suggest that a quiescence-like adaptation represents a generalizable stress tolerance strategy. These results raise intriguing questions about whether similar physiological state transitions contribute to microbial persistence in other kinds of fluctuating environments [28] and whether such mechanisms can be harnessed for industrial and medical applications, including bio-preservation and cryo-tolerance. By demonstrating that cells can evolve resilience via a physiologically mediated mechanism that operates alongside genetic factors, our work expands the understanding of adaptive evolution and stress biology beyond classical mutation-driven models [29].

## RESULTS

### A. *Saccharomyces cerevisiae* adapts to freeze-thaw stress

We first establish an experimental protocol in order to perform the selection (Methods; Figure 1**A**). It is known that the survival of *Saccharomyces cerevisiae* upon freeze-thaw depends on the rate of freezing and thawing. Survival has been found to be minimal when it undergoes a fast freeze (*>* 10 K/min) to −196^◦^C [30]. We chose this regime to work in to possibly achieve the most significant increase in survival after adaptation. Before proceeding, we determined the effect of duration at −196^◦^C on survival by exposing the Wild Type - CEN.PK a/ *α* (WT) stationary phase cell cultures to a fast freeze (*>* 10K/min) to −196^◦^C and allowed them to remain at −196^◦^C for varying lengths of time before thawing. Freezing was done by immersing 500 *µ*l of stationary phase culture in 1.5 mL tubes in liquid nitrogen (−196^◦^C) and the cultures were then thawed by immersing them in 25^◦^C water for 15 minutes. We note that the survival fraction of yeast does not vary with duration at −196^◦^C (Supplementary Information). Therefore, for experimental convenience, we used a freeze duration of 15 minutes and selected for freeze-thaw tolerance at high cooling rates (*>* 10 K/min).

**FIG. 1.**
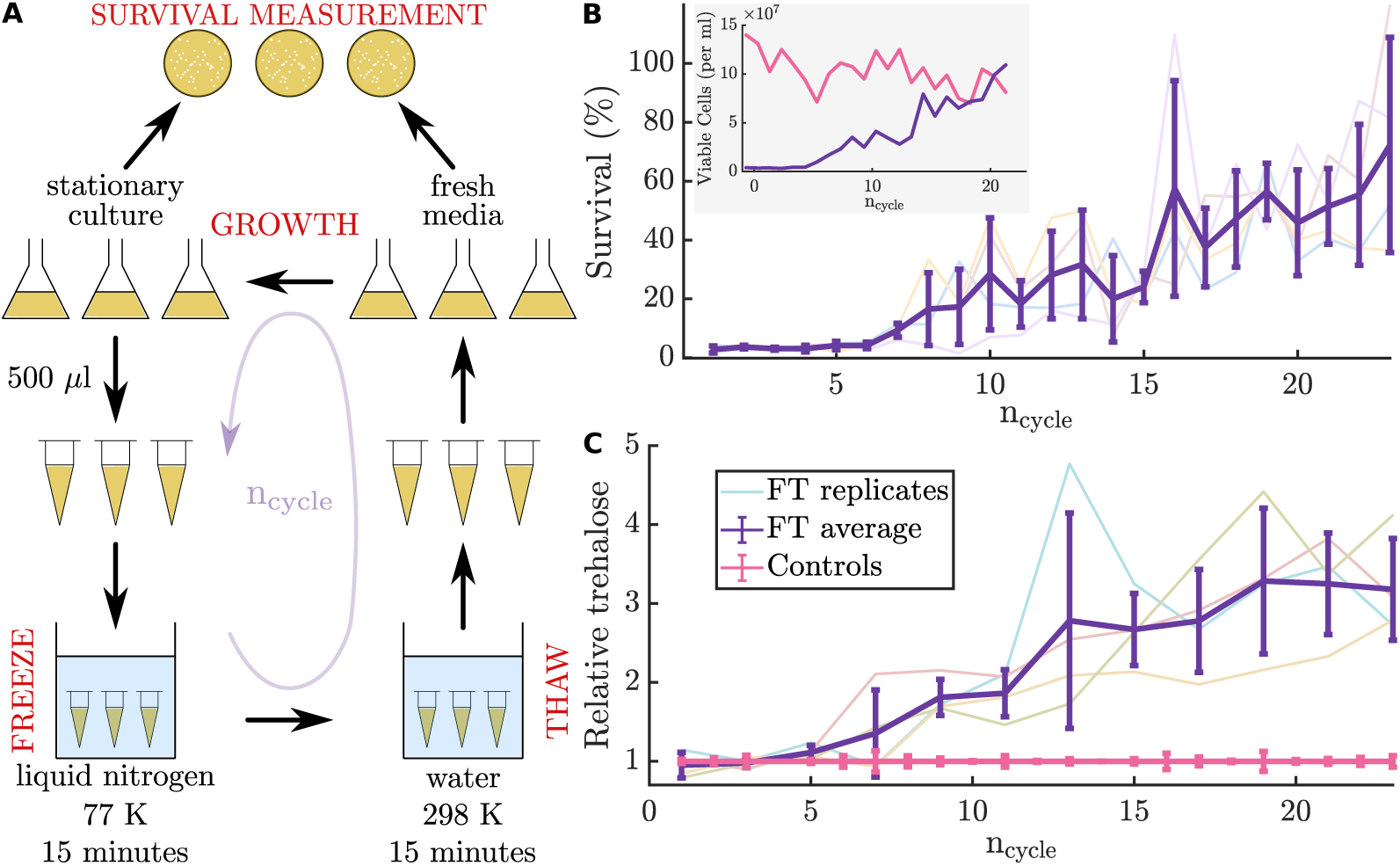
Experimental evolution can be used to increase survival of yeast under freeze-thaw stress. **A** Experimental design of selection for freeze-thaw tolerance. **B** Survival fraction of cells after freeze-thaw over the course of evolution. Each point on the thick purple line is the average of four biological replicates and two technical replicates for each biological replicate. Error bars indicate standard deviation. Each light coloured line represents a biological replicate. The evolution was performed thrice and the data from one evolution experiment is shown here. Inset: Average number of viable cells after every freeze-thaw-growth cycle. Purple - evolving population. Pink - control population. **C** Basal intracellular concentrations of trehalose relative to the control measured over the course of evolution. Each point on the thick purple line is the average of 4 biological replicates and 12 technical replicates for each biological replicate. Error bars indicate standard deviation. Each light coloured line represents a biological replicate. This measurement was made during one evolution experiment.

To start the experiments, a single yeast colony was picked from an agar plate and grown as liquid culture to stationary phase. This starter culture was divided into six parts — two for controls and four biological replicates for selection. The four biological replicates underwent freeze-thaw and the survival fraction was then measured by colony forming units (CFUs). The four populations that underwent freeze-thaw were independently grown in fresh medium until stationary phase (24 hours, ≈4 generations) and were independently put through freeze-thaw again. This cycle of freeze-thaw followed by growth was repeated several times until the survival fraction stopped increasing. For controls, the same protocol was followed without the freeze-thaw procedure.

Under this selection protocol, the survival fraction increased from ≈2% up to an average of ≈70%, with one of the replicates as high as 85%, in only a span of ≈25 freeze-thaw cycles. This shows that the yeast cells adapt quite rapidly to freeze-thaw in this regime of very low initial survival rates (i.e. ≈2%) and the dynamics are remarkably similar across independent evolution lines (Figure 1**B**). Next, we characterized the differences in the evolved strain with respect to the WT.

### B. Freeze-thaw tolerant populations of cells have higher basal intracellular levels of trehalose

Trehalose is a sugar containing two *α*-glucose units with a 1,1-glycosidic bond between them. Several studies have found a strong correlation between intracellular trehalose concentration and survival after freeze-thaw [31, 32]. Chen *et al.* showed that trehalose was sufficient to confer freeze-thaw tolerance when cooled slowly to −80^◦^C [33]. These studies either measure the amount of trehalose in existing freeze-thaw tolerant strains or the survival in trehalose accumulated mutant strains. Motivated by these studies, we measured the amounts of intracellular trehalose at the stationary phase to calculate the basal levels of trehalose present in cells, over the course of adaptation to freeze-thaw. Cell colonies were collected and weighed from plates used to measure survival using CFUs 48 hours after thawing. The concentration of trehalose was then measured using a previously described protocol [34] (Methods). We found that the amount of basal intracellular trehalose systematically increased over the course of adaptation to freeze-thaw, exhibiting similar dynamics to those of the cellular survival levels (Figure 1**C**). Basal levels of trehalose increased nearly 3 fold over the course of the selection.

Previous studies suggest that trehalose might play a role in reducing membrane damage [35], which has been implicated as the primary cause of death after freeze-thaw [36–38]. Therefore, we next turned our attention to investigating if the freeze-thaw tolerant cells have a membrane-damage related adaptation.

### C. Freeze-thaw tolerant cells undergo reduced membrane damage

The presence or absence of membrane damage was measured by staining the cells with two dyes — 5-carboxyfluorescein diacetate (5-CFDA) and Propidium Iodide (PI) — and measuring the intensities of fluorescence of both dyes using flow cytometry (Methods). The fluorescent form of CFDA leaks out of a cell quickly if it is membrane damaged. PI permeates cells with damaged membranes. Taken together, these dyes allow a clear distinction between membrane-damaged cells and cells with intact membranes, showing up as two distinct clusters in the data (Figure 2**A,B**). Each point on these plots represents a single cell. We compute a single metric that combines this information to estimate the total membrane damage (Methods), which clearly indicates that a smaller fraction of freeze-thaw tolerant cells undergoes membrane damage after freeze-thaw (Figure 2**C**). Further, we sought to understand the effect of membrane damage on cellular survival. We used fluorescence-activated cell sorting (FACS) to sort the cells into “membrane-damaged” and “membrane-intact” fractions. We then measured the number of colony forming units (CFUs) within each sorted fraction to quantify their viability (detailed description in Supplementary Information). We found that the survival fraction of freeze-thaw tolerant cells within the membrane-damaged sub-population exceeded the wild type by about two orders of magnitude (Figure 2**D**). Similarly, the viability of freeze-thaw tolerant cells with intact membranes was an order of magnitude higher than the corresponding wild type population.

**FIG. 2.**
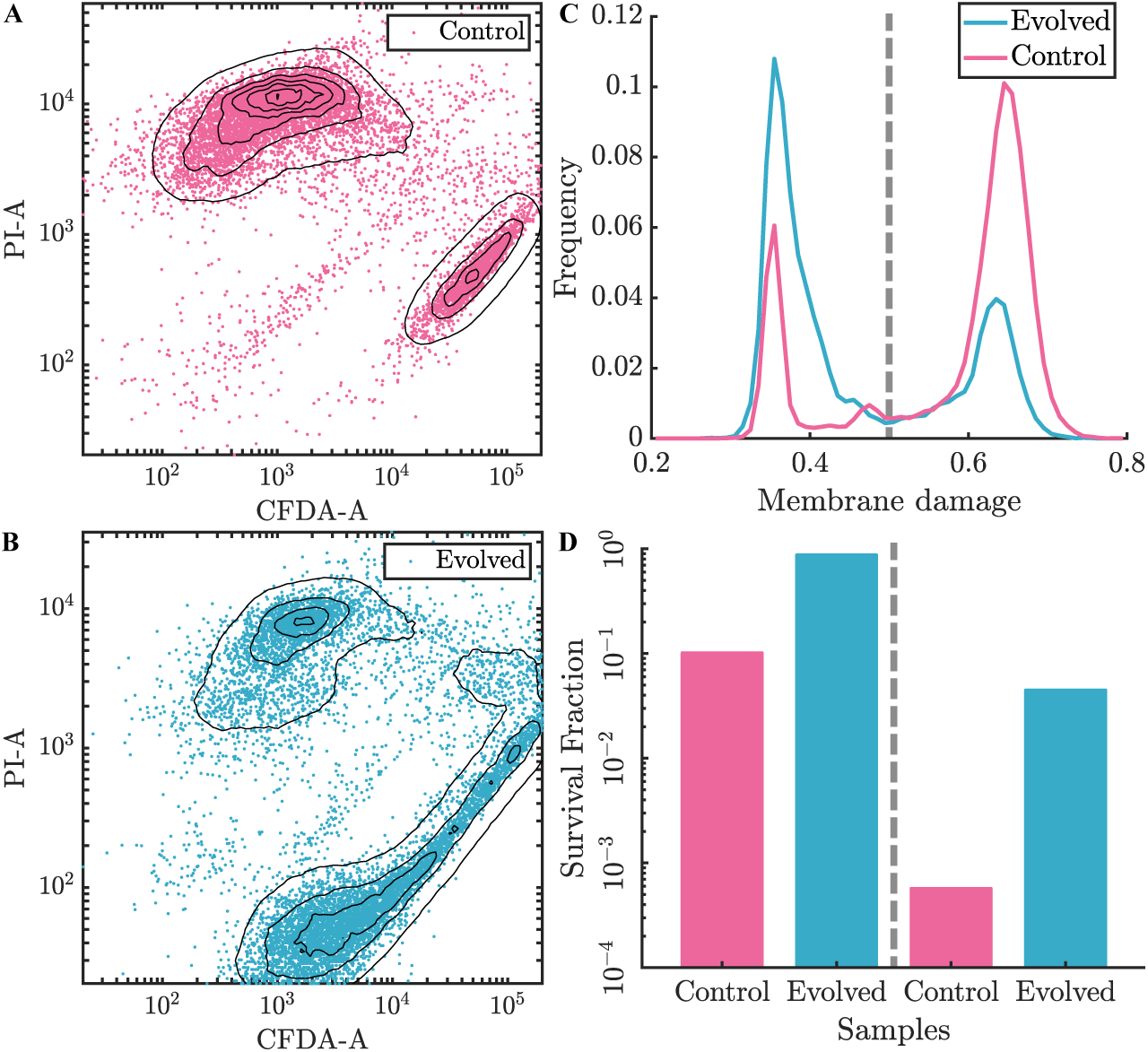
Freeze-thaw tolerant cells undergo reduced membrane damage and survive membrane damage. **A** Flow cytometry measurements on control cells. Each point represents a single cell and the plot contains data for 10000 cells. The x-axis is the fluorescence intensity in the 5-CFDA channel. The y-axis is the fluorescence intensity in the propidium iodide channel. The contour lines represent changes in density of cells, starting from 17 with increments of 17. The cluster on top comprises membrane damaged cells and the cluster at the bottom comprises cells with intact membranes. **B** Flow cytometry measurements on freeze-thaw tolerant cells. Each point represents a single cell and the plot contains data for 10000 cells. The x-axis is the fluorescence intensity in the 5-CFDA channel. The y-axis is the fluorescence intensity in the propidium iodide channel. The contour lines represent changes in density of cells, starting from 17 with increments of 17. The cluster on top comprises membrane damaged cells and the cluster at the bottom comprises cells with intact membranes. **C** The 2-dimensional data from **A** and **B** are used to compute a 1-dimensional distribution of “membrane damage” in cells. Area under the normalized curves: Evolved (Membrane intact) = 0.6347, Evolved (Membrane damaged) = 0.3127, Control (Membrane intact) = 0.2331, Control (Membrane damaged) = 0.7597. **D** Viability of cells measured after sorting the two clusters identified from the data in **A** and **B**. A 0.3% sorting error was measured by sorting beads. The entire experiment described in this figure was performed four times and the data from one experiment is shown here.

These data indicate that: (i) a significant fraction of cellular death following freeze-thaw is unrelated to membrane damage and (ii) the evolved cells have mechanisms that can aid in survival following membrane damage. This suggests cellular mechanisms and attributes beyond just the membrane protective roles of trehalose.

Increased intracellular trehalose is a defining characteristic of cellular quiescence — a temporary non-proliferating state — during which cells are tolerant to a range of stress conditions [39]. Therefore, we next investigated if the freeze-thaw tolerant cells exhibit traits similar to those of quiescent cells [40]. We started by probing the morphological and physical traits of freeze-thaw tolerant cells in the stationary phase – the point in their population growth stage at which the freeze-thaw stress was imposed.

### D. Freeze-thaw tolerant cells are smaller, denser, and exhibit reduced budding

Quiescent cells have been shown to be smaller and denser /[41]. Therefore, to investigate if there could be any morphological changes associated with the increased survival of cells, we imaged cells of the wild-type yeast and the freeze-thaw tolerant yeast at stationary phase to measure their size (Figure 3**A**). We observed a two-fold reduction in cell area (Figure 3**B**), which corresponds to an estimated six-fold volume reduction in freeze-tolerant cells. A decrease in cell volume could be due to a decrease in its mass and/or increase in cell density. By measuring the single cell mass density of cells at stationary phase using a density gradient column, we found that the freeze-thaw tolerant cells had a significantly higher cell density (Figure 3**C**).

**FIG. 3.**
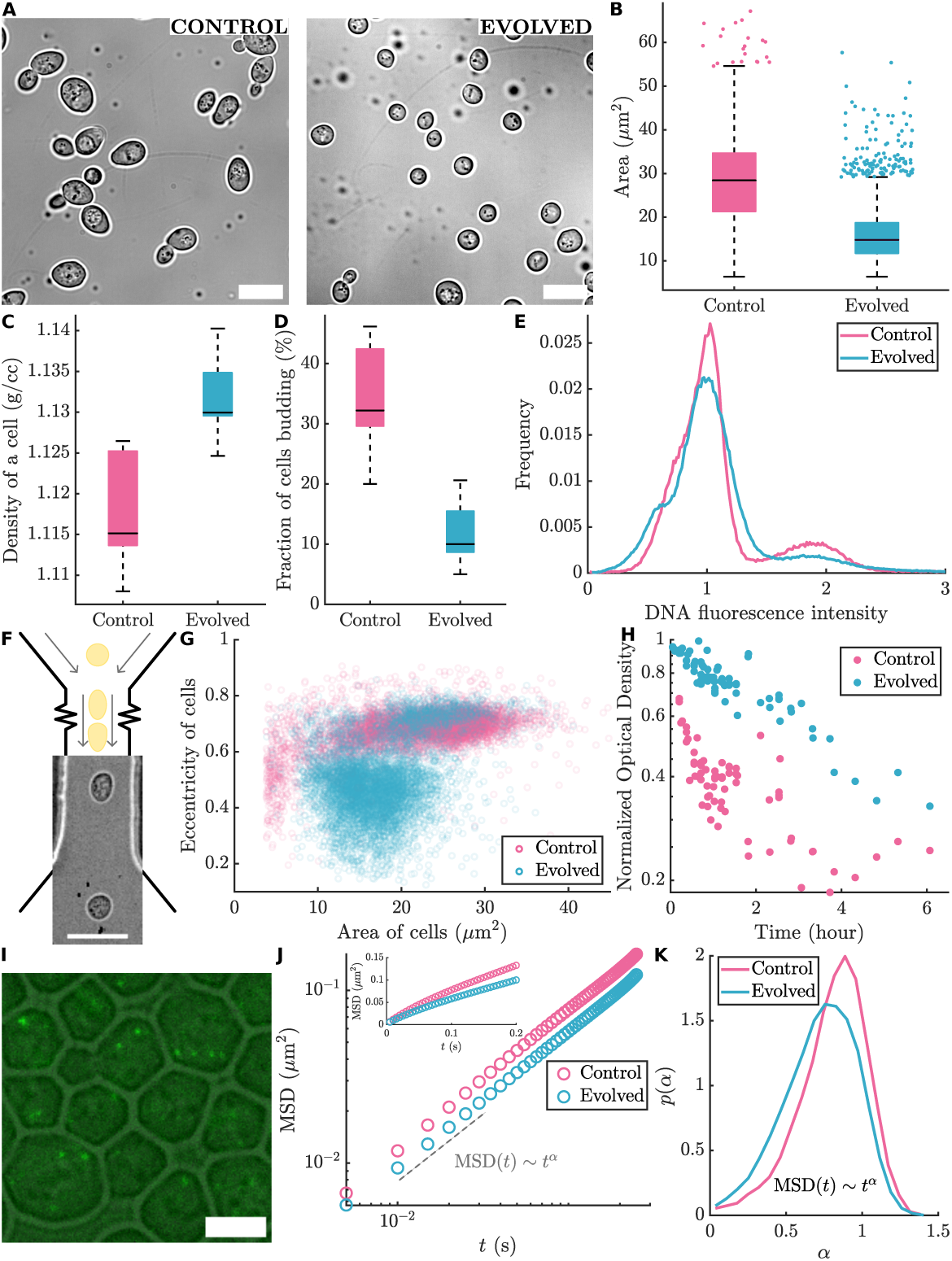
Cells of the population adapted to survive freeze and thaw are smaller, denser and exhibit reduced budding. **A** Brightfield image of cells. Scale bar - 10*µ*m. **B** Projected area of cells measured from images as shown in A. On each box, the central mark indicates the median, and the bottom and top edges of the box indicate the 25th and 75th percentiles, respectively. The whiskers extend to the most extreme data points not considered outliers, and the outliers are plotted individually as points scattered in the x-direction. Sample size (number of cells measured): Control = 1107, Evolved = 4281, Welch’s *t*-test: *P* = 1.3 *×* 10^−213^. This experiment was performed twice and the data from one experiment for one evolved population is shown here. **C** Single cell mass density measured using a percoll gradient column. On each box, the central mark indicates the median, and the bottom and top edges of the box indicate the 25th and 75th percentiles, respectively. The whiskers extend to the most extreme data points not considered outliers (there are no outliers). Welch’s *t*-test: *P* = 3.2 *×* 10^−7^. This experiment was performed twice and the data from both experiments for eight technical replicates of one biological replicate was pooled. **D** Fraction of cells budding in stationary phase measured from images as shown in A. On each box, the central mark indicates the median, and the bottom and top edges of the box indicate the 25th and 75th percentiles, respectively. The whiskers extend to the most extreme data points not considered outliers (there are no outliers). Sample size (number of cells measured): Control = 309, Evolved = 422, Welch’s *t*-test: *P* = 2.59 *×* 10^−6^. This experiment was performed twice and the data from one experiment for one evolved population is shown here. **E** Distribution of normalized fluorescence intensities of DNA in cells stained with SYTOX Green measured using flow cytometry. Area under the normalized curves: Evolved (Single DNA intensity) = 0.8726, Evolved (Double DNA intensity) = 0.1274, Control (Single DNA intensity) = 0.8330, Control (Double DNA intensity) = 0.1670. Sample size (number of cells measured): Control = 199682, Evolved = 399113. This experiment was performed twice and the pooled data for four biological replicates from one experiment is shown here. **F** Design of the microfluidic channel through which cells are flowed to measure deformation with an image of cells leaving the microfluidic channel. Scale bar - 15*µ*m. **G** Eccentricity and area of cells measured in the narrow channel. Flow rate: Control - 5.5cm/s, Evolved - 5.6cm/s. Sample size (number of cells measured): Control = 5825, Evolved = 6192. This experiment was performed four times for one evolved population and the data from one experiment is shown here. **H** Fraction of intact cells over time measured by turbidity of cells placed in water after Zymolyase treatment. This experiment was performed five times for one evolved population and three technical replicates and the pooled data for all five experiments is shown here. **I** Image of Genetically Encoded Multimers (GEMs) in the cytoplasm of cells. Scale bar - 5*µ*m. **J** Mean squared displacement (MSD) of GEMs tracked in the cytoplasm of cells, plotted on logarithmic axes. Inset: MSD plotted on linear axes. More than 10,000 trajectories were used to calculate the average. **K** Distribution of the exponent of time, *α*. More than 10,000 trajectories were used to plot the histogram.

From the cellular morphologies, we also found a reduction in the incidence of budding in the evolved cell population (Figure 3**D**), suggesting that they stopped doubling before reaching stationary phase and could have exited the proliferating phase of the cell cycle - a sign of quiescence entry. This is also corroborated by a measurement of total amount of cellular DNA — the evolved cell population has reduced fraction of dual DNA copies (Figure 3**E**), suggestive of quiescence.

The increased cellular mass density suggests that there could be an associated increase in molecular crowding in the cytoplasm, thereby reducing cytoplasmic mobility and increasing cellular stiffness. Since we also observed reduced membrane damage, we investigated if the freeze-thaw tolerant cells are more mechanically resilient.

### E. Freeze-thaw tolerant cells are mechanically stiffer

Yeast cells are naturally ellipsoid due to the presence of a stiff cell wall. Exposure to Zymolyase, a cell-wall digesting enzyme, leads, over time, to the formation “spheroplasts”, spherical cells bound by a lipid membrane. We used these Zymolyase treated cells to quantify their stiffness, by employing deformability cytometry a method previously described by Otto *et al.* [42]. Briefly, the cytometry is performed by suspending the Zymolyase treated cells in a viscous buffer and flowing them through a narrow microfluidic channel. The resultant shear causes cellular deformation, which is measured using high-speed imaging (Figure 3**F**). The deformation of cells due to shear is quantified by their eccentricity in relation to their size. For a given cell size, lower cell stiffness results in a higher deformation and, therefore, higher eccentricity. From such a measurement, we found two clusters of freeze-thaw tolerant cells with a majority of cells minimally deformed, compared to the freeze-thaw sensitive (WT) cells (Figure 3**G**). The presence of two clusters indicates increased cellular stiffness even after the long Zymolyase treatment. Munder *et al.* found that cells that had increased cytoplasmic stiffness did not relax into spheres after treatment with Zymolyase [19]. Hence, the relaxation of freeze-thaw tolerant cells from ellipsoidal to spherical upon treatment with Zymolyase could be affected either by their cytoplasmic stiffness or their cell wall resilience due to a varied cell wall composition that is not affected by Zymolyase to the same extent. Therefore, we further investigated the role of these two factors – cell wall resilience and cytoplasmic stiffness – in increased cellular stiffness.

In yeast, the cell wall maintains the structure of the cell. When the cell wall is removed and cells are placed in a hypo-osmotic environment, cells lyse as they cannot regulate the amount of water permeating the cell. Therefore, to investigate the presence of the cell wall after digestion with Zymolyase, we exposed cells to Zymolyase for varying lengths of time in an appropriate buffer followed by a hypo-osmotic environment *i.e.* pure water (Supplementary Information). Cells lacking cell-wall lyse in water. So, we measured the turbidity of the solution after placing the cells in water as a measure of the fraction of intact cells. A significant fraction of the freeze-thaw tolerant cells do not lyse even after exposure to Zymolyase for long periods of time, compared to the duration taken for most of the freeze-thaw sensitive cells to lyse (Figure 3**H**). This suggests three possibilities - (a) the freeze-thaw tolerant cells have an altered cell wall composition that is not easily digested by Zymolyase, (b) the cells have an altered membrane structure (as seen above) that does not allow water to enter the cell even after cell wall digestion, or (c) an increased cytoplasmic stiffness that does not allow cells to relax after cell wall digestion and hence the cellular components do not spread into solution enough to change the turbidity of the solution.

These results suggest that cells of the adapted population have increased mechanical (structural) resilience associated with modified cell wall or cytoplasmic properties. Next, we specifically investigated the cytoplasmic rheology to understand the nature of the increased cellular stiffness.

### F. Freeze-thaw tolerant cells exhibit reduced cytoplasmic mobility

To study the cytoplasmic mobility, we tracked the intracellular motion of engineered fluorescent particles (41 nm diameter) (Figure 3**I**). Briefly, to produce these fluorescent particles, we transformed the wild type CEN.PK strain to express genetically encoded multimers (GEMs) using a plasmid developed by Delarue *et al.* [43]. Delarue *et al.* replaced the INO4 promoter with the monomeric fluorophore sequence fused with the INO4 promoter and selected for the integrants using uracil auxotrophy. We inserted the monomeric fluorophore fused with the INO4 promoter at the HO locus in a prototroph which does not affect growth characteristics significantly in a diploid strain [44]. The plasmid also contained a KanMX marker for selecting the integrants using resistance to G418 (full details in the Supplementary Information). Evolution was performed on the wild type strain transformed to express GEMs. We imaged the GEMs expressed in the cells using total internal reflection fluorescence (TIRF) microscopy, used image analysis to track the particles and analysed the trajectory dynamics using custom written MATLAB code.

To quantify the mobility of these particles, we calculated the mean squared displacement (MSD) for several trajectories of many cells. Two-dimensional MSD is related to the time window, *t*, in which it is calculated by MSD(*t*) ∼ *t^α^*, where the exponent, *α*, is related to the dynamics of the particles. We calculated the MSD across all trajectories of all replicates, with each line on the plot representing an average of 10’s of thousands of single particle trajectories. We found that the MSD is lower for particles in the cytoplasm of freeze-thaw tolerant cells (Figure 3**J**). This suggests reduced mobility of particles in the cytoplasm of freeze-thaw tolerant cells. From the time and ensemble-averaged MSD, we find *α* ≈ 0.83 for particles in the cytoplasm of freeze-thaw sensitive cells and *α* ≈ 0.77 for particles in the cytoplasm of freeze-thaw tolerant cells. The probability density function, *p*(*α*), of the exponent, calculated by fitting the time-averaged MSD of a particle (Supplementary Information) to a power law, further reveals a larger fraction of particles exhibiting a significantly restricted motion in the freeze-thaw tolerant cells compared to the control cells (Figure 3**K**). We find decreased cytoplasmic mobility and increased confinement of particles in the cytoplasm across all freeze-thaw adapted replicate lines (more details in the Supplementary Information).

### G. Freeze-thaw tolerant cells exhibit quiescence-like properties

The morphological and physical attributes that we described thus-far have been associated with the entry of yeast cells into a quiescent state [40]. It is known that trehalose promotes efficient cellular entry into quiescence and re-entry into the cell cycle [39]. Further, higher intracellular trehalose causes a decrease in growth rate of the cell upon re-entry into exponential phase [45, 46]. Higher trehalose levels also lead to a decrease in lag phase duration [16, 47–49]. Therefore, we measured the population growth characteristics of the control and evolved populations after a freeze-thaw at stationary phase. The optical density (OD) of the cell cultures (absorbance at 600nm) was measured to obtain the relative cell concentrations over the course of growth using a spectrophotometer (Figure 4**A**).

**FIG. 4.**
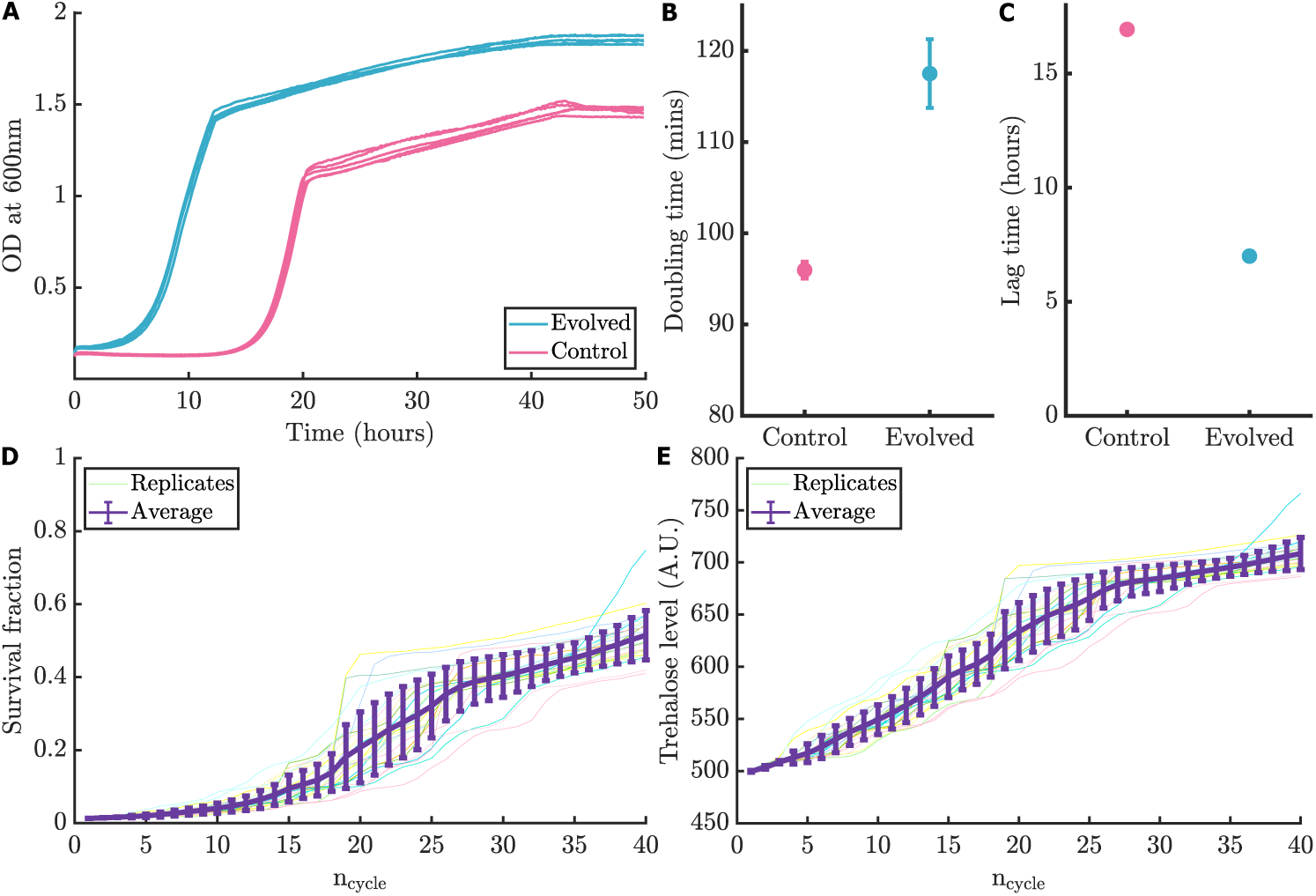
Selection based on trehalose linked entry into a quiescence-like state recapitulates the adaptation dynamics. **A** Optical density of the cell cultures inoculated into fresh media after a freeze-thaw. **B** Average doubling time of cells in the population calculated from the growth curves in A. **C** Average lag phase duration of cells in the population calculated from the growth curves in A. **D** Survival fraction of cells after freeze-thaw simulated over the course of evolution. Each light coloured line represents an iteration of the simulation. The thick purple line indicates the average for 25 iterations. Error bars indicate standard deviation. **E** Basal intracellular concentrations of trehalose relative to the control simulated over the course of evolution. Each light coloured line represents the average of the population in an iteration of the simulation. The thick purple line indicates the average for 25 iterations. Error bars indicate standard deviation of 25 iterations.

The growth rates of the populations were calculated for the glucose metabolism phase of exponential growth from the maximal slope of the OD curve. Indeed, we found that the minimum doubling time for the evolved populations was longer than that for the control populations (Figure 4**B**). Further, the control population of cells exhibited a 60% longer lag duration (calculated as time taken to reach the maximal growth rate) than the evolved population of cells (Figure 4**C**). Therefore, we observe co-evolution of a freeze-thaw tolerant, trehalose-rich state together with rapid regrowth under a stress-growth alternating regime. These results corroborate previous studies that associate higher intracellular trehalose levels with decreased growth rate and lag phase duration and, therefore, transition to a quiescent state.

Based on these data, we hypothesized that the freeze-thaw tolerant population is enriched with quiescent cells which leads to a high survival fraction. In this case, selection for quiescence would be equivalent to selection for freeze-thaw tolerance. However, it is not feasible to experimentally select for quiescence; therefore, to investigate this hypothesis, we numerically simulated the adaptation of yeast based on the ability of cells to enter and exit quiescence.

### H. Selection based on trehalose linked entry into a quiescence-like state recapitulates the adaptation dynamics

We developed a population level model with trehalose concentration acting as the sole phenotype under selection, since trehalose levels determine the growth rate, probability of entering quiescence, survival probability after a freeze-thaw, and lag phase duration before re-entry into the cell cycle [39] (full details of the model are in the Supplementary Information).

Briefly, we start with an initial cell population characterized by a Gaussian distributed phenotypic trehalose levels. The trehalose level for each cell in stationary phase is correlated with its growth rate in log phase, and assume that it directly determines (a) the probability of going into a quiescent state upon encountering starvation conditions [50, 51], (b) the probability of survival after freeze thaw [15, 52–55], and (c) the duration of lag phase upon re-entry into the cell cycle.

In each cycle of growth-freeze-thaw, the population has differing growth rates, and cells with lower trehalose values have a higher growth rate and a larger number of offspring. Each offspring is allowed a “mutation” which acts to shift its stationary state trehalose concentration with respect to its parent. These mutational effects are drawn from a fat-tailed distribution symmetric about zero *i.e.* most mutations have no effect on the phenotype of the offspring, while the possibility of a few large effect mutations exists. Each cell division uses up a unit of a finite resource, and the population faces starvation when the resource runs out.

Upon starvation, the population differentiates into quiescent and non-quiescent cells. Cells with higher amounts of trehalose have a higher probability of entering a quiescent state. All non-quiescent cells are subsequently assumed to perish when the population is subjected to selection *i.e.* freeze-thaw. The survival of cells after freeze thaw is determined similarly, using a survival probability that increases with the intracellular trehalose concentration. The population that survives freeze-thaw is diluted by selecting a fraction of the population randomly, which is then allowed to enter growth again. The lag phase of each cell is further determined by its trehalose concentration, with cells with higher trehalose concentrations facing a shorter lag time.

Thus, the states of quiescence and increased survival under freeze thaw cause the population distribution to shift to higher values of mean trehalose. Similarly, cells with higher trehalose have longer periods of growth due to a shorter lag phase, and take over more of the population with each generation of offspring. This effect is diminished slightly by the fact that higher trehalose also leads to a smaller growth rate, but is not strong enough initially to stall the increase in mean trehalose of the population. At larger values of trehalose the effect of a decreasing growth rate becomes stronger, leading to saturation of trehalose levels over the course of evolution. We find that when cells are selected in this way for quiescence, the dynamics of the survival (Figure 4**D**) and trehalose levels (Figure 4**E**) of the adapting population qualitatively resemble the experimentally measured adaptation dynamics (Fig 1**B,C**). We speculate that the more sudden jumps seen in the experimental curves could be due to the larger number of replicates in the simulation than what the experiments than what allow for. Altogether, our results suggest that cellular entry into a quiescence-like state facilitates the adaptation to freeze-thaw.

### I. The mechano-chemical adaptation is convergent

Finally, we ask if the results we describe thus far are a result of the specific selection protocol we used for experimental evolution or if they point to a more general mechanism. Therefore, we sought to investigate the phenotypes emerging from a different freeze-thaw selection regime.

To do this, we developed a selection protocol based on exposing cells to multiple consecutive freeze-thaw cycles before a growth cycle (Figure 5**A**). First, we note that the survival fraction of the cells decreases exponentially with consecutive freeze-thaw exposures, reducing by about an order of magnitude with the number of consecutive exposures, k_FT_ (Figure 5**B**). Each point obtained by varying k_FT_ is, therefore, a quantitatively (and qualitatively) different selection regime. Although the results obtained so far have been obtained from the k_FT_ = 1 regime, we now report results from the k_FT_ = 3 selection regime. Interestingly, despite nearly two orders of magnitude reduction in the initial viability, the yeast adapted to the k_FT_ = 3 regime, with survival close to 40%. We find that the population of cells evolved in this regime also exhibits a multifold increase in basal trehalose levels (Figure 5**C**) and reduced budding fractions (Figure 5**D**). Furthermore, the MSDs obtained from cells of the evolved population in the k_FT_ = 3 regime fall directly on top of those obtained from the k_FT_ = 1 regime (Figure 5**E**); the distribution of the MSD exponents, *p*(*α*) is also quite similar (Figure 5**F**). On the basis of these findings, we conclude that the mechano-chemical phenotype promoting cellular entry into a quiescence-like state is a convergent adaptation.

**FIG. 5.**
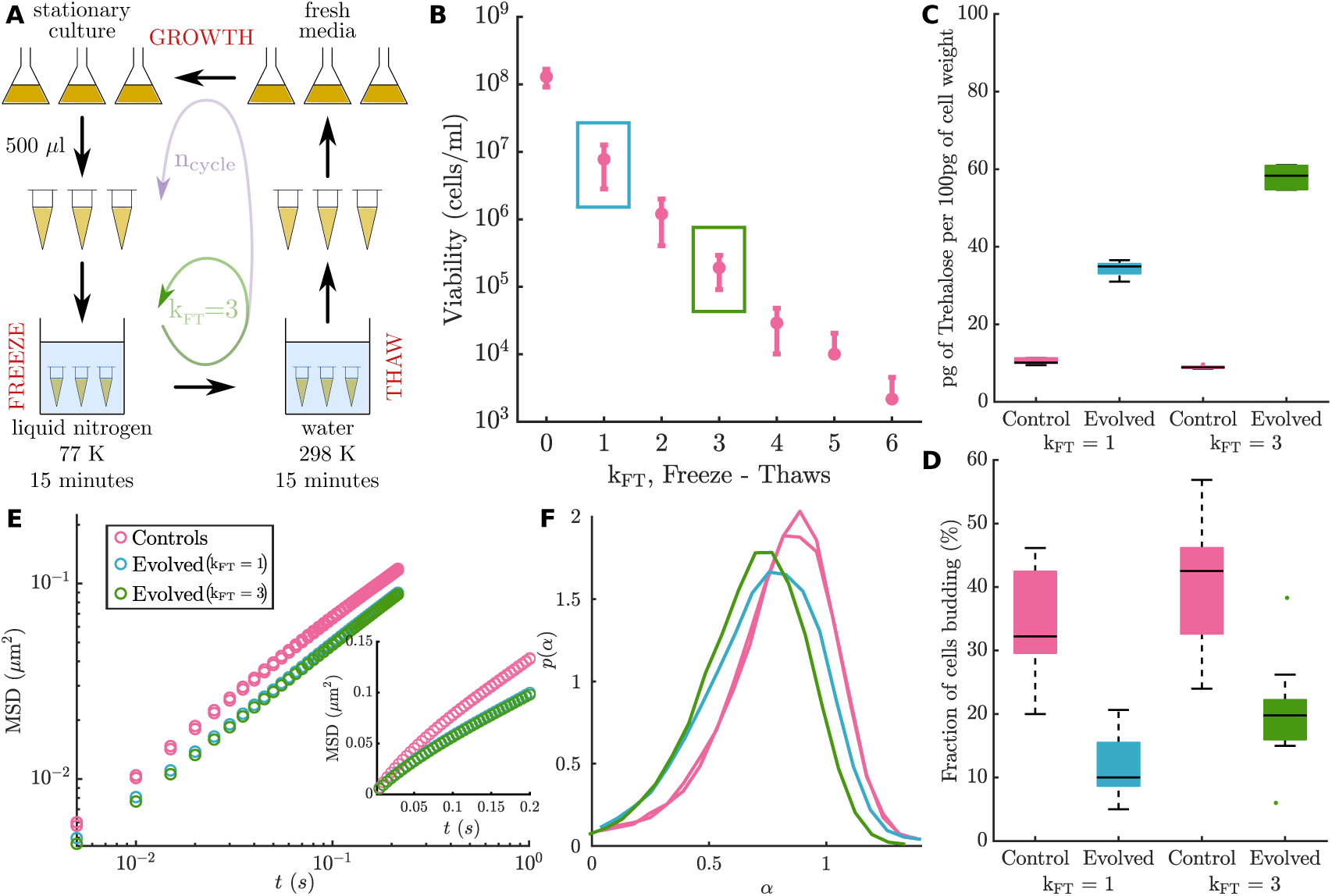
The mechano-chemical adaptation to freeze-thaw tolerance is convergent. **A** Experimental design to measure survival after refreezing and for the evolution of refreezing tolerance. **B** Number of viable cells after consecutive freeze-thaws. Error bars indicate standard deviation. This experiment was performed five times with two technical replicates and the pooled data from all experiments is shown here. The point marked in blue denotes the selection pressure for (*k_F T_* = 1) evolution. The point marked in green denotes the selection pressure for (*k_F T_* = 3) evolution. Basal amounts of intracellular trehalose measured in the stationary phase. On each box, the central mark indicates the median, and the bottom and top edges of the box indicate the 25th and 75th percentiles, respectively. The whiskers extend to the most extreme data points not considered outliers (there are no outliers). *P* (*k_F T_* = 1) = 7.1 *×*10^−8^, *P* (*k_F T_* = 3) = 1.6 *×*10^−7^. **D**Fraction of cells budding measured in the stationary phase. On each box, the central mark indicates the median, and the bottom and top edges of the box indicate the 25th and 75th percentiles, respectively. The whiskers extend to the most extreme data points not considered outliers, and the outliers are plotted individually as points scattered in the x-direction. This experiment was performed once and the data for one evolved population and six technical replicates is shown here. Sample size: Control (*k_F T_* = 1) = 309, Evolved (*k_F T_* = 1) = 422, *P* (*k_F T_* = 1) = 2.5 *×* 10^−6^, Control (*k_F T_* = 3) = 357, Evolved (*k_F T_* = 3) = 459, *P* (*k_F T_* = 3) = 8.1 *×* 10^−5^. **E** Mean squared displacement (MSD) of GEMs tracked in the cytoplasm of cells, plotted on logarithmic axes. Inset: MSD plotted on linear axes. More than 10,000 trajectories were used to calculate the average. **F** Distribution of the exponent of time, *α*. More than 10,000 trajectories were used to plot the histogram.

### J. Distinct genetic routes underlie the adaptations

Since we found a convergent phenotype across all replicates of different freeze-thaw regimes, we sought to understand if the phenotype is a physiological response with epigenetic inheritance or whether it was more likely to be genetically inherited. We propagated the evolved lines without freeze-thaw stress for more than 25 generations and found that survival fraction did not reduce significantly (details in the Supplementary Information), suggesting genetic inheritance. We then performed whole genome sequencing to investigate if the adaptations are associated with a particular genotype (or a specific genetic program).

DNA from cells of samples of the evolved and control populations, collected from the middle and end of the evolution experiment, along with the parent strain was isolated. Sequencing was performed on whole individual populations for the evolved and control replicates and from a single clone for the parent strain. Since we were interested in shared and high-frequency genetic routes, we performed population sequencing even though it might result in under-sampling of rare variants. The sequenced reads were trimmed and filtered based on quality and then mapped to a reference genome. The mapped reads were analyzed post-filtering for the frequency of occurrences of Single Nucleotide Polymorphisms (SNPs), insertions, and deletions in each replicate population. Further, we also analyzed the genome for copy number variation. We developed a model to estimate the frequency of copy number variants in a population based on the coverage for each gene for each sample (full details in the Supplementary Information). The final list of all such genetic variants obtained for the freeze-thaw tolerant populations, along with their function, are shown in Table V. Each variant is marked for its mutation type and the sample(s) in which it was detected.

We then checked if any variants were present across multiple samples, as this would imply that the mechano-chemical phenotype found across all replicates was a consequence of a specific genotype(s). We found no variants common across all replicates of both k_FT_ = 1 and k_FT_ = 3 selection regimes. It must be noted that mutations in different genes with similar or shared functions, or genes linked in cellular pathways, could lead to the same adaptive phenotype. We investigated this aspect using the curated information available on the *Saccharomyces* genome database (SGD) on regulatory, metabolic, and physical interactions. We sought to identify a common molecular function or pathway that could have been targeted by different variants across samples. Therefore, we looked for interactions between the different genes and checked if any of the genes are regulators of other genes in the list to understand if there is any complementarity in the genotypes found such that certain replicates have related genotypes. However, no such known cellular network could be found. We also considered changes in gene-copy numbers as a potential basis for observed phenotypes, particularly the increase in intracellular levels of trehalose. Again, we found no gene with a consistent copy-number variation across all replicates (details in the Supplementary Information). These results reveal that the convergent mechano-chemical adaptations seen in this evolution experiment are achieved through multiple non-overlapping genetic routes.

## DISCUSSION

Using experimental evolution, we have shown that *Saccharomyces cerevisiae* can adapt to high freeze rate perturbations in a stereotypical manner, with survival increasing nearly two orders of magnitude over 25 cycles of freeze-thaw-growth. In all our replicate lines of evolution, the basal intracellular level of trehalose, a sugar speculated to play a physical role in maintaining membrane and cytoplasmic integrity [23, 56, 57], exhibited a significant (three-fold) increase in the adapted cells. The adapted cells also show smaller sizes, higher mass densities, and reduced membrane damage. Using rheological measurements we showed that the adapted cells are stiffer and had lower particle mobility within the cytoplasm. Whole genome sequencing indicated that the increased survival is associated with multiple, distinct genotypes.

Earlier, independent studies have pointed out the role of increased trehalose to confer stress tolerance in yeast, while not connecting the increased trehalose to a broader physiological context (the trehalose was increased artificially via extracellular means) [58]. Studies have also associated cytoplasmic stiffening with transition to dormancy/quiescence in yeast [19, 59] while not looking at such physical changes in an evolutionarily adaptive sense. Further, Sleight et al used experimental evolution with bacteria to shown that microbes can indeed evolve to increase survival to freeze-thaw stress – in this case, the survival increased only 2 fold, compared to the nearly 40 fold in our case – but concluded that “the physiological mechanism for this fitness improvement remains unknown” [29]. To our knowledge, our work is the first study showing the *de novo* linkage between increased intracellular trehalose, reduced membrane damage, decreased cytoplasmic mobility, increased cellular stiffness and transition to a quiescence-like state.

**TABLE I.** A listing of the genetic mutations across all replicates including both the k_FT_ = 1 and k_FT_ = 3 selection regimes, coloured boxes indicate the identified mutations. Replicates of populations of cells adapted to freeze-thaw stress do not have any specific genetic mutation in common. FS - Frame Shift, SC - Stop Codon, NS - Non-Synonymous, S - Synonymous, IG - Inter-genic region.

| Gene | Function | $k_{FT} = 1$<br>Rep 1 | $k_{FT} = 1$<br>Rep 2 | $k_{FT} = 1$<br>Rep 3 | $k_{FT} = 1$<br>Rep 4 | $k_{FT} = 3$<br>Rep 1 | $k_{FT} = 3$<br>Rep 2 | $k_{FT} = 3$<br>Rep 3 | $k_{FT} = 3$<br>Rep 4 |
| --- | --- | --- | --- | --- | --- | --- | --- | --- | --- |
| Cln3 | Cyclin (cell cycle regulator) |  |  |  |  |  |  | S |  |
| Pho3 | Phosphate metabolism |  |  |  |  |  |  |  | S |
| Ho | Homothallic switching endonuclease |  |  | NS | NS |  |  |  | NS |
| SCEN_L05230 | Hypothetical protein | NS |  |  |  |  |  |  |  |
| Itr1/Itr2 | Myo-inositol transporter | NS |  |  |  |  |  |  |  |
| Bch2/Chs6 | Chitin Synthase-related |  |  |  | NS |  |  |  |  |
| Vac8 | Vacuolar inheritance |  |  | FS | FS |  |  |  |  |
| Ade5,7 | De novo purine nucleotide pathway |  |  | NS |  |  |  |  | NS |
| SCEN_J00260 | Hypothetical protein |  |  | IG |  |  |  |  |  |
| Alr1 | Aluminum Resistance |  |  |  | NS |  |  |  |  |
| Dnf1/Dnf2 | Aminophospholipid translocase |  | NS |  |  |  |  |  |  |
| Pma1 | Plasma Membrane ATPase |  | NS |  | NS |  |  |  |  |
| Yeh1/Yeh2 | Yeast sterol ester hydrolase |  |  |  | IG |  |  |  |  |
| SCEN_L03720 | Hypothetical protein |  |  |  |  |  |  |  | NS |
| Cbk1 | Cell wall biosynthesis kinase |  |  |  |  |  | SC |  |  |
| Alg8 | Asparagine-linked glycosylation |  |  |  | S |  |  |  |  |
| Ras1/Ras2 | Adenylate cyclase activating GTPase | NS | NS |  |  | NS | NS | NS |  |
| Rpc82 | RNA polymerase C subunit |  |  |  |  | NS |  |  |  |

The high levels of trehalose could play multiple roles in the increased mechanical resilience of the cell. Reduced membrane damage in freeze-thaw tolerant cells could be a result of altered lipid composition, altered cell wall composition and increased trehalose. Several studies show that increased membrane fluidity of cells was associated with higher freeze-thaw tolerance, with trehalose hypothesized to aid in membrane fluidization [29, 36, 60]. In addition, our results on the increased cellular stiffness suggest that the cell wall composition could also be altered, thereby protecting the cell wall associated membrane from undergoing damage.

A major cause of death after freeze-thaw has been suggested to be ice crystal formation [23, 26, 61, 62]. Ice crystals can damage the cell membrane, proteins in the cytoplasm and disrupt the intracellular structure and organisation, thereby hindering the revival of cellular processes upon thawing. A glassy cytoplasm with decreased mobility is a cytoplasmic rheological state that could aid in retaining cellular structure and organisation during freeze-thaw transitions. Indeed such a cytoplasmic state has been implicated in response to various environmental stresses [19, 20, 43], also to be a role of trehalose [57] and has long been associated with freeze-thaw tolerance. We found the *de novo* encoding of a similar cytoplasmic state with increased intracellular trehalose in the adapted freeze-thaw tolerant cells.

Trehalose has been shown to be a good glass-former, compared to other sugars like glucose and sucrose [63] and has also been found to localise in the cytosol [55]. Several other studies shown that higher intracellular trehalose levels are associated with tolerance to desiccation, heat, ethanol etc [58, 64]. Cakar *et al.* allowed mutagenic lines of *Saccharomyces cerevisiae* to adapt to various kinds of stresses and found that freeze-thaw tolerant populations were tolerant to multiple other kinds of stress such as high temperature, ethanol and oxidative stress [65]. The evolution of a mechano-chemically reinforced survival phenotype underscores the role of biophysical constraints in shaping cellular adaptation. Cytoplasmic stiffening and increased trehalose accumulation likely serve as protective mechanisms against mechanical and osmotic damage [19, 20]. This highlights how stress tolerance is not solely dictated by genetic changes but also by modulations in intracellular material properties [13]. Beyond yeast, similar adaptations have been observed in bacterial biofilms and dormant cancer cells [66–68], suggesting broader implications for microbial resilience, antibiotic tolerance, and tumor dormancy.

Interestingly, our sequencing analysis did not reveal any mutations directly related to trehalose synthesis, degradation or transport (Table V). Therefore, this increased concentration of trehalose is not of any direct genetic origin, unlike previous studies that used trehalose synthesis, degradation or transport mutants and related viability after freeze-thaw to intracellular trehalose concentration [31–33].

While freeze-thaw is lethal to most kinds of cells, several cells survive freeze-thaw on ecological and evolutionary timescales. There have been a few previous studies that used experimental evolution to obtain freeze-thaw tolerance in cells. Kwon *et al.* started with a *L. rhamnosus* strain that had a 60% survival fraction after freeze-thaw and the adapted strain was found to have a ≈1.5 fold increase in survival fraction over 150 freeze-thaw-growth cycles [60]. Sleight *et al.* found only a 2.5 times increase in survival fraction of *E. coli* over 150 freeze-thaw-growth cycles (1000 generations) [29]. Although, these studies are on very different species, they both find *cls* mutants that increase membrane fluidity to be associated with survival. Since these kinds of stress do not cause a drastic reduction in survival, it could lead to a more gradual phenotypic change as fitness is not affected as much – it must be noted that in our experiments, the survival of wild type *Saccharomyces cerevisiae* to rapid freeze-thaw is a few percent or lesser. By using freeze-thaw regimes where survival is affected significantly, we found that *Saccharomyces cerevisiae* adapts to have orders of magnitude higher survival in only ≈25 freeze-thaw-growth cycles (≈100 generations). Our observations of stark phenotypic changes could be because of the strong selection regime in which we operate.

The various cellular attributes that we have discovered in our study correlate with those of quiescent cells. The molecular mechanisms of quiescence entry and exit and the characteristics of the quiescent state are only recently being understood [41, 47] — specifically it has been uncovered that trehalose promotes efficient entry into and exit from quiescence. Our simulations on trehalose-mediated selection for quiescence suggest that an entry to a quiescence-like state, as well as the ease of re-entering the growth cycle (through lag) facilitates the adaptation dynamics we see in the experiments.

Several studies have found that quiescent cells are more stress resilient [69], although the underlying mechanisms underlying have not yet been understood. Specifically, the genetic basis for quiescence is not fully explored. While certain genes involved in the stress response leading to quiescence have been studied through enrichment of quiescent cells in the population [70], these genetic mutations are not present among the adapted populations in our experiments. The emergence of a quiescence-like state in our experimental evolution study, suggests that increased mechanical resilience of a quiescent cell, rather than any specific genetic basis, could be responsible for stress resilience.

By bridging experimental evolution, biophysical characterization, and predictive modeling, our work provides a framework for investigating how cells can rapidly acquire stress tolerance through state transitions rather than relying solely on genetic mutations. Future research should explore the molecular regulation of these adaptive states, including potential epigenetic contributions and metabolic shifts that facilitate production of high trehalose and quiescence entry and maintenance under extreme conditions [71].

## ACKNOWLEDGMENTS

We thank Sunil Laxman for discussions and help with the project. We acknowledge funding and support from the Max Planck Society through a Max-Planck-Partner-Group and also the Department of Atomic Energy (India), under project no. RTI4006, and the Simons Foundation (Grant No. 287975). We thank the Central Imaging and Flow Cytometry Facility, Sequencing Facility and computational facility at NCBS. The authors also acknowledge the Sane Lab (NCBS) for access to a high speed camera and the Mayor Lab (NCBS) for access to TIRF microscopy.

## SUPPLEMENTARY INFORMATION

### Strains and growth

The yeast strain *Saccharomyces cerevisiae* CEN.PK a/*α* obtained from Laxman lab (InStem) was used as wild type to evolve increased tolerance to freeze thaw. Cultures were grown in rich nutrient media called YPD (Yeast extract [1%] (HIMEDIA, Catalog no. - RM027), Peptone [2%] (HIMEDIA, Catalog no. - RM014), Dextrose [2%] (Qualigens, Catalog no. - Q15405), with 2% agar (HIMEDIA, Catalog no. - GRM026) for plates).

### Freeze-thaw

500 *µ*l of stationary phase culture was aliquoted into 1.5 ml microcentrifuge tubes. Freezing was done by immersing the tubes in liquid nitrogen for 15 minutes, unless stated otherwise. The culture was then thawed by removing the tubes from liquid nitrogen and immersing them in room temperature water for 15 minutes.

### Selection to freeze thaw

Figure 1 shows the protocol followed to select for adaptation to freeze-thaw stress. Yeast cultures were grown for 24 hours (≈12 generations) to attain stationary phase. The culture was then put through either one freeze-thaw or three freeze-thaws according to the experiment. The cells that survived the freeze-thaw(s) were grown in fresh media and the cycle of freeze-thaw(s)-growth was repeated until the survival levels saturated. For the controls, the same protocol was followed without the freeze-thaw procedure.

### Survival measurements

Survival in cells/ml was measured for each cycle during the evolution by checking for Colony Forming Units (CFUs) on a YPD agar plate. The concentration of viable cells was measured before and after the freeze-thaw(s) to estimate the fraction of cells that survived the freeze-thaw(s). Cell culture was appropriately diluted and added to a Petri dish with YPD agar poured into it along with 4-7 glass beads. The culture was then spread on the plate by shaking the petri dish and the beads were discarded. The petri dishes were then incubated in a 30^◦^ C incubator for 48 hours. The petri dishes were then imaged and the number of cell colonies formed were counted.

**FIG. S1.**
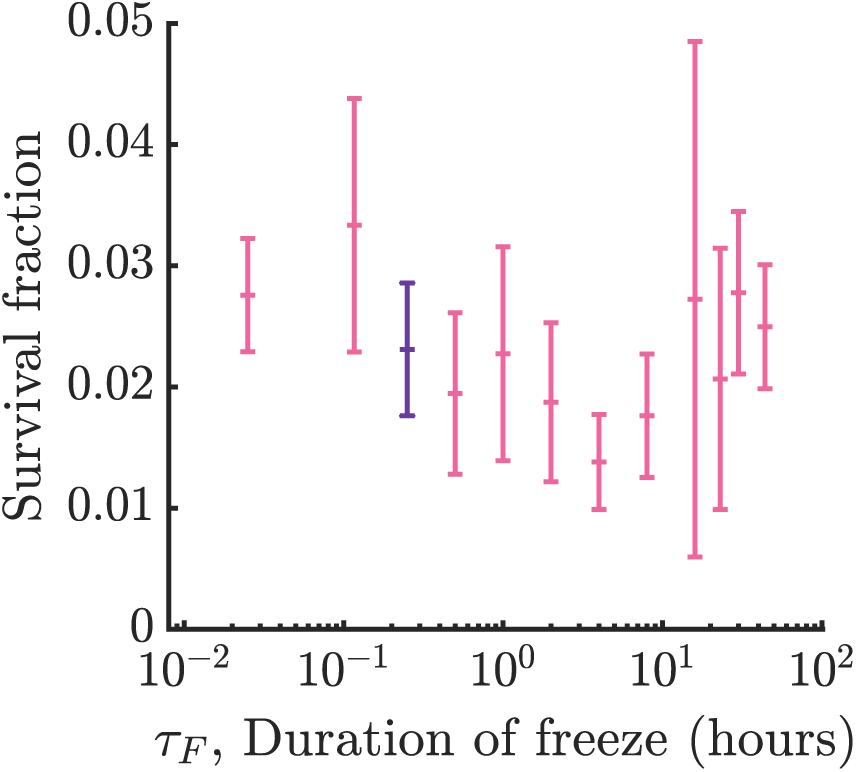
Survival fraction of cells measured over different freeze durations Error bars represent standard deviation of biological and technical replicates.

### Intracellular trehalose measurements

Intracellular levels of trehalose were measured based on a previously described protocol [34]. During evolution of freeze-thaw tolerance, cell colonies were collected in microcentrifuge tubes from plates used to measure CFU 48 hours after thawing. Cells were weighed and suspended in 0.25 ml of 0.25M sodium carbonate. For measurements after the evolution experiment, 500ul of stationary phase culture was centrifuged at 3000 rcf for 2 minutes to remove media. The cell pellet was then washed with Phosphate Buffered Saline and suspended in 0.25 ml of 0.25M sodium carbonate.

The tubes were gently sonicated for a minute and then incubated at 95^◦^C for 4 hours to lyse the cells and break down all existing glucose. The suspension was buffered to pH 5.2 by adding 0.15 ml of 1M acetic acid and 0.6 ml of 0.2M sodium acetate for trehalase activity. The mixture was then incubated with 0.025U/ml trehalase (Sigma-Aldrich, Catalog no. - T8778) at 37^◦^C overnight with shaking to break down trehalose into two glucose molecules. The suspension was then centrifuged at 16,900g for 5 minutes and the supernatant was assayed for glucose using a glucose assay kit (Sigma-Aldrich, Catalog no. - GAGO20). The glucose assay was performed using a 96-well plate. 40 *µ*l of the appropriately diluted supernatant was added to each well so that the glucose concentration was within the dynamic range of the assay (20-80 mg/ml). The plate was incubated for 5 minutes at 37^◦^ C, followed by the addition of 80ul of the assay reagent to each well to start the colorimetric reaction. The plate was then incubated at 37^◦^ C and the reaction was stopped after 30 minutes by adding 80ul of 12N sulfuric acid to each well. The absorbance of each well was measured at 540 nm to calculate the concentration of glucose (and therefore trehalose) from a standard curve.

### Membrane damage and survival measurements

Lysis of cells after freeze-thaw was checked using BD FACSVerse™ Cell Analyzer by measuring the concentration of intact cells before and after freeze-thaw. To investigate membrane damage, cells were stained using the FungaLight™ Yeast CFDA, AM/Propidium Iodide Vitality Kit (Invitrogen, Catalog no. - F34953). Cells were then analysed using a BD FACSAria™ Fusion Flow Cytometer. The cells clustered into two populations on the CFDA vs propidium iodide plot - one membrane-damaged population and one membrane intact population. The two populations were sorted and the concentration of cells in each population was analysed on the BD FACSVerse™ Cell Analyzer. Further, the sorted cell cultures were treated with 50 ug/ml Ampicillin and spread on YPD agar plates with appropriate dilutions to measure survival by counting the number of Colony Forming Units (CFUs). We also checked that FACS did not affect the viability of the resultant fractions. A membrane damage metric was obtained from the fluorescence intensities of propidium iodide (PI) and 5-CFDA (CFDA) combined as follows:

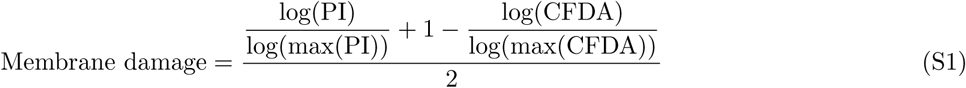

### Cell morphology measurements

Cells were imaged at stationary phase to measure their size and shape. Brightfield imaging on cells was performed using an inverted Olympus IX-81 microscope equipped with a Photometrics Prime sCMOS camera, a motorized stage, 100x/1.4NA UPlanSApo oil immersion objective and controlled with OLYMPUS cellSens Dimension software. The area and eccentricity of cells was then measured from these images using custom written MATLAB code. The fraction of cells budding was manually counted from these images.

### Single cell mass density measurements

Single cell mass density was measured using a Percoll gradient column. Stock Isotonic Percoll (SIP) was made by adding 1.5M sodium chloride to Percoll™ (GE Healthcare life sciences, Catalog no. - 17-0891-01) in a 1:9 ratio. SIP was diluted with 0.15M sodium chloride to make 70%, 80% and 90% density solutions. 1.5 ml of each of these solutions was aliquoted in different 2 ml microcentrifuge tubes. The tubes were centrifuged at 20,000g at 4^◦^ C for 15 minutes to prepare density gradient columns. 500 *µ*l of stationary phase cells were washed once in Phosphate Buffered Saline (PBS) by centrifugation at 200g for 2 minutes and resuspended in 100 *µ*l PBS. Density was calibrated by using a mixture of Density Marker Beads (Cospheric, DMB-kit). After the density gradient columns were prepared, 100 *µ*l of cell and bead suspensions were gently added at the top of the columns ad centrifuged for 1 hour at 400g at room temperature. The columns were then imaged using a phone’s camera and the position of cells was measured to calculate their density.

### DNA fluorescence intensity measurements

Cells were stained with SYTOX Green based on a previous protocol [72]. 3ml of stationary phase culture was centrifuged at 1500g for 2 minutes at room temperature to remove media. The cells were fixed in 70% ethanol and incubated for 16 hours at 4^◦^ C. Cells were then pelleted and washed in 50mM sodium citrate buffer (pH 7.5). The cell pellet was re-suspended in 500 *µ*l RNAse A buffer (500ul sodium citrate buffer + 5ul RNAse A) and incubated at 50^◦^ C for 2 hours with shaking. Cells were then washed and resuspended in 50mM sodium citrate and incubated at 30^◦^ C for 3 hours with shaking. Cells were pelleted and resuspended in Proteinase K solution (20ul enzyme (2mg/ml) + 180ul sodium citrate buffer). They were incubated for 37^◦^ C for 1 hour, with shaking. Following Proteinase K treatment, cells were pelleted, washed and resuspended in 1 ml of sodium citrate buffer and incubated overnight at 30^◦^ C with shaking. The sample was diluted 20 times and 1 drop of SYTOX Green was added to 500ul of the sample. The samples were then analysed using flow cytometry. The SYTOX green fluorescence intensity was acquired through the FITC channel.

### Deformability cytometry

Deformability cytometry was performed on cells with a protocol based on a previous study by Otto *et al.* [42].

**FIG. S2.**
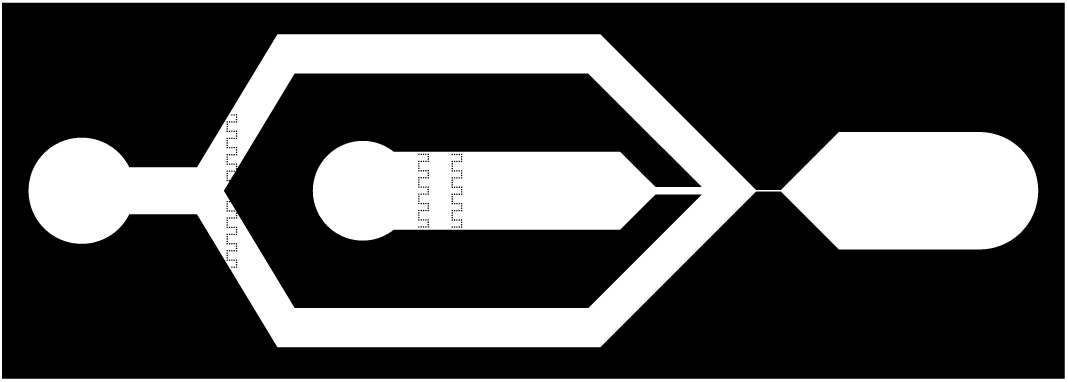
Design of the deformability cytometry device

#### a. Microfluidic Device Preparation

The microfluidic device was made using polydimethylsiloxane (PDMS) (Sylgard 184; Dow Corning) from a silicon wafer mould and bonded to a coverglass. The silicon wafer mould was fabricated using a soft lithography protocol in a clean room from a quartz mask with design as shown in S2. The following steps detail the protocol for fabrication of the silicon wafer mould:

1. A silicon wafer was washed with acetone, ethanol and isopropyl alcohol and air dried.
2. It was then placed on a hot plate at 95^◦^ C for 20 minutes for all the solvent to evaporate.
3. The wafer was plasma cleaned for 10 minutes.
4. The wafer was spin coated with SU 8 5 with the following parameters:
5. The wafer was pre-softbaked at 65^◦^ C for 3 minutes.
6. It was then softbaked at 95^◦^ C for 12 minutes.
7. The quartz mask was placed on the silicon wafer in a mask aligner and exposed to UV light for 3 seconds using vacuum contact.
8. Pre post exposure bake was done at 65^◦^ C for 1 minute.
9. Post exposure bake was done at 95^◦^ C for 4 minutes.
10. The silicon wafer was washed with developer for 3 minutes.
11. It was then washed with isopropyl alcohol and acetone
12. The wafer was finally dried on a hot plate at 95 ^◦^ C.
13. The features were measured on a profilometer.

**TABLE II.** Parameters for spin coating

| Cycle | Hold (Time (s)) | Speed (rpm) | Acceleration |
| --- | --- | --- | --- |
| 1 | 5 | 500 | 200 |
| 2 | 30 | 875 | 300 |

The following steps elaborate the protocol used for preparation of PDMS devices from the silicon wafer:

1. PDMS and the cross linker were mixed thoroughly in a 10:1 ratio and poured over the silicon wafer mould.
2. The PDMS was placed in a desiccator for degassing until it was completely free of bubbles.
3. PDMS was cured at 70^◦^ C for two hours.
4. The device was cut out and holes were punched for inserting tubing at the inlet and outlet.
5. The PDMS was cleaned with scotch tape. A coverglass was cleaned with Hellmanex, ethanol and isopropyl alcohol and air dried.
6. The PDMS and coverglass were plasma cleaned for 3 minutes.
7. The PDMS was covalently bonded to the coverglass by placing the two in contact with each other and pressing lightly.
8. The microfluidic device was then placed at 70^◦^ C for 10 minutes to ensure complete bonding.

#### b. Solutions

- 0.5% 2-hydroxyethyl cellulose (Sigma Aldrich, Catalog no. - 434981) + 1.2M Sorbitol in PBS (buffer for cells and sheath fluid)
- 1.2M Sorbitol in PBS (SB, for spheroplasting)
- 1.2M Sorbitol + 1% Glucose in PBS (SB+Glc, for spheroplasting)
- 100% 2-mercaptoethanol (BME, for spheroplasting)

#### c. Material

- Two 250uL Glass Syringes (Hamilton, 1725)
- Syringe Pump (Harvard Apparatus, Pump33 DDS)
- Two Needles (Dispovan 1.2 x 38 mm)
- Three pieces of silicon tubing (OD 1.65 mm, Dow Corning Silastic)
- Deformability cytometry microfluidic device

#### d. Viscous Buffer Preparation Procedure

*T*he following ingredients were used for preparation of the viscous buffer used to shear cells:

- 100 mL Autoclaved Phosphate Buffered Saline (Sigma Aldrich, Catalog no. - P3813)
- 0.5g 2-Hydroxyethyl cellulose, with a molecular weight around 1.3 × 10^6^ (Sigma Aldrich, Catalog no. - 434981)
- 21.8604g D-Sorbitol (Sigma Aldrich, Catalog no. - S1876)

### Procedure

1. 0.5g of 2-Hydroxyethyl Cellulose was added to 100 mL of autoclaved PBS.
2. The mixture was stirred at 65°C until the cellulose dispersed homogeneously. This process, lasting about 2 hours, resulted in increased viscosity and a slight turbidity.
3. 21.8604g of Sorbitol was added to obtain a concentration of 1.2M. Stirring at room temperature continued for a few hours until the solution turned clear.
4. The solution was filtered using a 0.22*µ*m Merck syringe filter and left at room temperature.

#### e. Cell Spheroplasting

- 1 ml of stationary phase cell culture was centrifuged at 1500g for 1.5 minutes.
- The cell pellet underwent two washes with Sorbitol in PBS (SB) by centrifuging at 1500rcf for 1.5 minutes.
- The washed cells were resuspended in 1 mL of SB with the addition of 2 L of 100% 2-mercaptoethanol (BME).
- The cells were then incubated at 30 ^◦^ C for 45 minutes.
- After BME treatment, cells underwent further washing (3 times with Sorbitol + 1% Glucose in PBS (SB+Glc)) by centrifuging at 1000rcf for 2 minutes.
- A mixture of 1 mL SB+Glc and 100 L of 2.5 mg/mL Zymolyase was added to the cells, and the suspension was incubated at 30°C.
- The WT cells were incubated for 1 hour, while the EVO cells were incubated for 4 hours.
- The resulting spheroplasts were washed twice with SB by centrifuging at 1000rcf for 2 minutes.
- Spheroplasts were resuspended in the prepared buffer.
- The suspension was subjected to vortexing for 2 minutes to break up the clumps.
- A bumpy surface, such as a pipette tip rack, was employed for additional clump disruption.
- The suspension was degassed in a desiccator for 1 minute to eliminate air bubbles.

### Microfluidic Device Setup

- Two clean and dry glass syringes were wiped with 70% ethanol.
- Silicon tubing was connected to the syringes using needles.
- The tubing was cleaned with an air gun and the free ends of the tubing were wiped with 70% ethanol.
- The sheath syringe and its tubing were filled with filtered buffer, ensuring that the tubing was submerged in buffer to avoid air bubbles.
- The cell syringe and its tubing were filled with spheroplasts in buffer, avoiding air bubbles.
- Sheath tubing, followed by cell tubing and discard tubing, were carefully inserted into the designated holes of the microfluidic device, using coverage of other holes with Scotch tape to prevent debris ingress.

#### f. Imaging and Flow setup

- Images were acquired on an Olympus IX-81 inverted microscope with a Phantom Miro X4 high-speed camera using a 60X oil objective. The camera was controlled using Phantom Camera Control (PCC) 3.4 software.
- Syringes were connected to the syringe pump and the free end of the discard tube was placed in a microcentrifuge tube secured to the stage.
- The inlets were filled with sheath fluid and cells. The rest of the device was then filled with sheath fluid.
- Then, the sheath fluid was set to flow at the specified rate followed by cells.
- The camera was focused to capture cells as they exited the channel, ensuring a consistent region of interest (ROI).
- The following camera parameters were used:

**–** Resolution = 128 x 64
**–** FPS = 40k
**–** Exposure = 4us
**–** Brightness = 0
**–** Gain = 0.73
**–** Gamma = 0.95
**–** Sensitivity = 2.5
- The system was monitored until the flow stabilized, as indicated by a constant number of frames for a cell to travel a fixed distance.
- A waiting time of 10-15 minutes was necessary for flow stabilization.
- Movies were stored as .cine files for fast acquisition.

#### g. Analysis

The .cine files were first converted to smaller files in TIFF format using the PCC software such that the first frame was empty to use as background image for segmentation. Image segmentation and data analysis were performed using custom written code on MATLAB.

### Kinetics of cell disintegration

Cell culture in stationary phase was centrifuged at 1500g for two minutes and the supernatant was removed. Half the initial volume of Sorbitol Buffer was added and cells were treated with 0.2% 2-mercaptoethanol for 45 minutes at 30^◦^ C with shaking. Cells were then washed three times with Enzyme Buffer by centrifuging at 1500g for two minutes. 0.25mg/ml Zymolyase was added to cells in the Enzyme Buffer and this reaction mixture was tested for spheroplasting. Cells in Enzyme Buffer was used as the control. 200ul of the reaction mixtures and controls were aliquoted into one set of wells of a 96-well plate. The plate was incubated at 30^◦^ C with shaking in a Spark® multimode plate reader throughout the experiment. At each time point, the plate was moved out, 30ul of reaction mixture (or control) and 150ul of water was added to a well and the absorbance was measured at 800nm.

### Cloning of GEMs gene into HO-poly-kanMX4-HO Plasmid Vector

Bases 3119 to 5315 of the *pRS305-Leu2-PINO4-Pfv-GS-Sapphire* plasmid (AddGene, Plasmid # 116930) were inserted into the *HO-poly-kanMX4-HO* plasmid vector (AddGene, Plasmid # 51662, obtained from Laxman lab, InStem). The region of the *HO-poly-KanMX4-HO* plasmid vector for cloning was determined based on the restriction enzyme sites BsaBI and BsiWI.

**FIG. S3.**
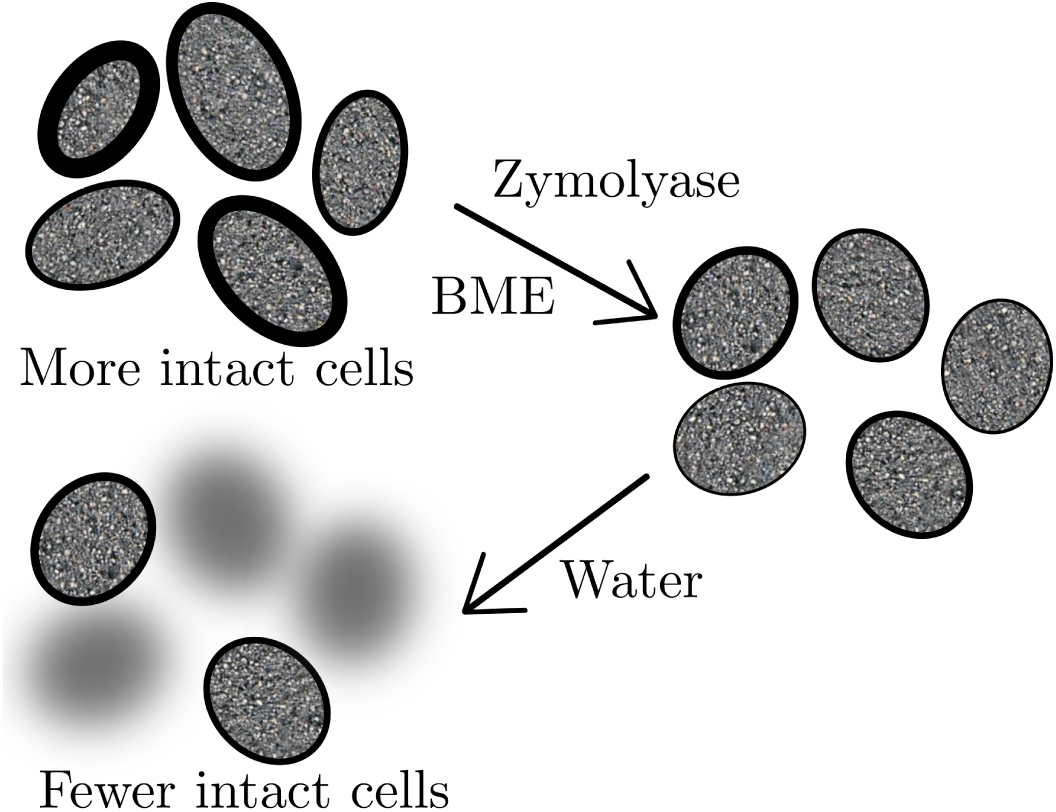
Sketch of the experimental design to probe the digestion of the cell wall. BME - beta-mercaptoethanol.

#### h. Primer Design and PCR Optimization

Primers were designed for PCR amplification of the yeGFP region and their parameters were tested using oligoanalyzer. The annealing temperature for the PCR was determined to be 62°C for template copying, with a first step annealing temperature of 55°C. Gradient PCR was performed to find the optimal annealing temperature.

#### i. Plasmid Isolation

Bacterial cultures containing the *HO-poly-KanMX4-HO* plasmid vector (obtained from Laxman lab, InStem) and the *pRS305-Leu2-PINO4-Pfv-GS-Sapphire* plasmid were inoculated into Luria Broth + Ampicillin (100 µg/ml) and incubated at 37°C. Plasmid isolation was performed using a MiniPrep kit (QIAGEN, Catalog no. - 27104). Plasmid quality and concentrations were measured using a TECAN NanoQuant plate reader.

#### j. PCR Amplification

The PCR amplification of the yeGFP region from the *pRS305-Leu2-PINO4-Pfv-GS-Sapphire* plasmid was performed using KOD hotstart DNA polymerase kit with the following components in the reaction mixture:

| Component | Final concentration | Volume |
| --- | --- | --- |
| 10X buffer | 1X | 5 µl |
| 25mM MgSO<sub>4</sub> | 1.5 mM | 3 µl |
| DNTPs (2 nM each) | 0.2 mM(each) | 5 µl |
| Sense primer (10 µM) | 0.3 µM | 1.5 µl |
| Anti-sense primer (10 µM) | 0.3 µM | 1.5 µl |
| Plasmid | 1ng/µl | 1 µl |
| Water |  | 32 µl |
| Polymerase (1 U/µl) | 0.02U/µl | 1 µl |
| **Total** |  | 50 µl |

Based on the 2200bp length of the fragment, the following parameters were used for the PCR:

1. Polymerase activation: 95^◦^ C for 2 minutes
2. Denaturation: 95^◦^C for 20s
3. Annealing: 54.3^◦^C for 30s
4. Extension: 70^◦^C for 45s
5. Repeated steps 2-4 for 35 cycles
6. Annealing remaining: 54.3^◦^C for 5 mins
7. Temperature was set to 4^◦^C.

**FIG. S4.**
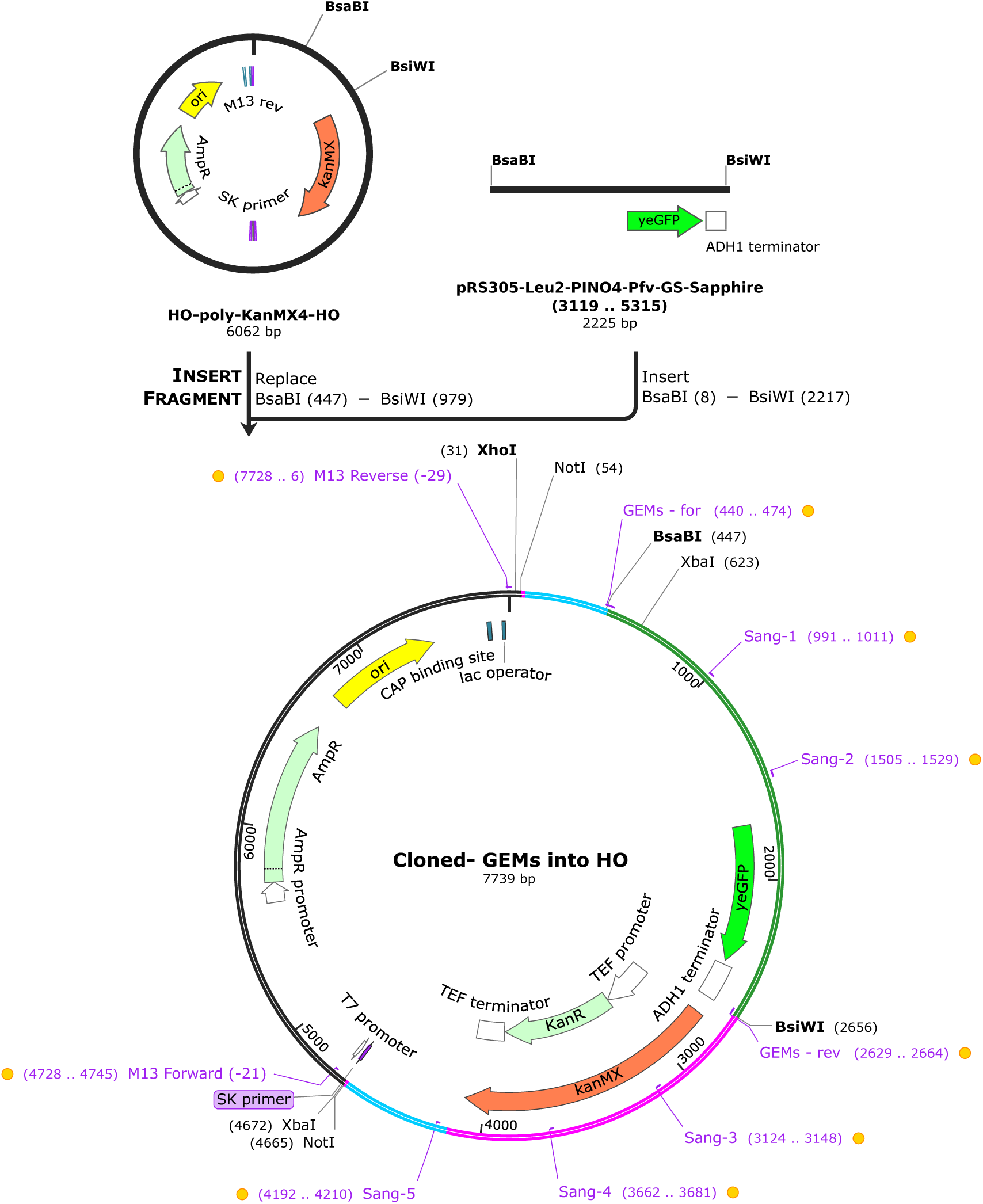
Map of cloned GEMs Plasmid

#### k. PCR Purification and Product Analysis

The PCR products were purified using a PCR purification kit (QIA-GEN, Catalog no. - 28104) to remove other reactants. Purified plasmid quality and concentrations were measured using a TECAN NanoQuant plate reader.

#### l. Restriction Enzyme Digestion

The purified PCR products and the *HO-poly-KanMX4-HO* plasmid vector were separately digested using BsiWI-HF and BsaBI-HF restriction enzymes. using the following protocol:

1. The following reaction mixtures were prepared:

PCR product (purified)

HO-poly-KanMX4-HO plasmid

|  |  |
| --- | --- |
| Nuclease free water | 84 μl |
| PCR product | 4 μl |
| CutSmart | 10 μl |
| BsiWI-HF | 2 μl |
| total | 100 μl |
| Nuclease free water | 82 μl |
| Plasmid | 6 μl |
| CutSmart | 10 μl |
| BsiWI-HF | 2 μl |
| total | 100 μl |

1. The reaction mixtures were incubated at 37^◦^C for 4:10 hours.
2. 2ul of BsaBI was added and the reaction mixtures were incubated at 60^◦^C for 3 hours.
3. The digested products were purified using a PCR purification kit (QIAGEN, Catalog no. - 28104) to remove restriction enzymes and other reactants.
4. The purified digested products were analysed using a TECAN NanoQuant plate reader.
5. The digested *HO-poly-KanMX4-HO* plasmid was run on a 1% agarose gel prepared with 50ml TAE and 2ul SYBR green dye at 100V for 3 hours alongwith an appropriate ladder.
6. The required 5.5kb band was cut out under UV.
7. The gel band was purified using a gel purification kit (QIAGEN, Catalog no. - 28704).
8. The purified product was analyzed on an agarose gel to confirm the presence of the desired DNA fragments.

#### m. Ligation

The insert was ligated into the vector using T4 DNA ligase (NEB). Different ratios of vector to insert were tested for ligation efficiency. The ligation mixtures were incubated at 16^◦^C overnight. The ligation mixtures were then heat inactivated by incubating at 65^◦^C for 10 minutes.

#### n. Transformation into Bacteria

The ligation mixtures were transformed into competent *DH5α* cells using the following protocol:

1. Competent cells at −80^◦^C were thawed on ice.
2. 20 *µ*l of ligation mixture was added to the competent cells and incubated on ice for 30 minutes.
3. A heat shock was provided by heating 42^◦^C for 90s and then keeping on ice for 90s.
4. 50ul of the cell suspension was added to 1 ml of LB.
5. Cells were incubated at 37^◦^C for 1 hour with shaking.
6. The transformed cells were plated onto LB+Kanamycin agar plates and incubated overnight at 37^◦^C.

Colonies from the transformed plates were picked and subjected to colony PCR using Taq pol and gel electrophoresis to confirm successful transformation.

#### o. Plasmid Isolation and Sequencing

Plasmids from positive colony PCR samples were isolated using the MiniPrep kit and submitted for Sanger sequencing to verify the sequence with primers as shown in S4.

#### p. Restriction Enzyme Digestion for Transformation

The plasmid was digested with NotI prior to transformation in order to be integrated into the yeast genome using the following protocol: The reaction mixture was incubated at 37^◦^C for 6 hours followed by 4^◦^C to stop the reaction.

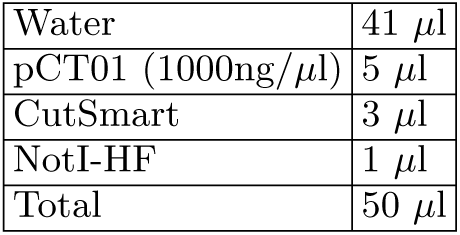

#### q. Yeast Transformation

1. 10ml yeast cell culture was grown to 1 OD.
2. A 1.0 ml sample of carrier DNA at a concentration of 2 mg/ml (salmon sperm) was boiled for 5 minutes and chilled in an ice/water bath. 1 mg/ml of carrier DNA was used in the reaction.
3. Yeast cells were spun down at 3000g for 1 minute.
4. The pellet was resuspended in approximately 1 ml of water and spun down again in a 1.5 ml microcentrifuge tube at 1000g for 1 minute.
5. The pellet was washed by resuspending in 800 *µ*l of 0.1 M Lithium Acetate (LiAc) and spinning down the cells at 1000g for 1 minute.
6. The following reagents were added to prepare the transformation mix:
7. PEG 3500 (50% w/v, sterile filtered): 240 *µ*l
8. Lithium Acetate (1.0 M): 35 *µ*l
9. Boiled (salmon sperm) carrier DNA: 25 *µ*l
10. 2 µg NotI digested plasmid + water: 50 *µ*l
11. 350 *µ*l of the transformation mix was added to the cells and vortexed vigorously.
12. Incubation was carried out at 30^◦^C for 30 minutes.
13. The tubes were then incubated in a 42^◦^C water bath for 30 minutes.
14. Centrifugation was performed at 1000g for 1 minute, and the transformation mix was removed.
15. 1.0 ml of YPD was pipetted into each tube; the pellet was stirred with a micropipette tip and vortexed.
16. Cells were spun down in the culture tube, media was removed, and they were resuspended in 100 *µ*l water. This 100 *µ*l was plated onto YPD agar plates + G418.
17. The plates were incubated at 30^◦^C until colonies appeared.

#### r. Colony Selection and Culturing

Transformed yeast colonies were selected, inoculated into YPD medium, and cultured overnight. A −80^◦^C stock of the culture was prepared for long-term storage.

### Particle tracking of GEMs

Single particle tracking of the GEM particles was performed on cells spread on an agar pad. The particles were imaged on a Nikon eclipse Ti microscope in partial TIRF mode using a 100x 1.45 NA Plan TIRF objective and a 488 excitation. Images were acquired every 5ms on a Photometrics prime 95B sCMOS camera. Particles were tracked and trajectories were analysed using custom written MATLAB code.

### Mean squared displacement analysis

The time-averaged mean squared displacement (MSD) for a single particle *i* in terms of its *x* and *y* coordinates is calculated as:

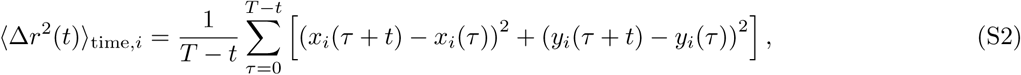

where:

- *x_i_*(*τ*) and *y_i_*(*τ*) are the *x*- and *y*-coordinates of particle *i* at time *τ*,
- *t* is the lag time,
- *T* is the total observation time for the trajectory of each particle.

This equation calculates the MSD for a single particle by averaging its squared displacements over all possible starting points *τ* for a given lag time *t*. The time and ensemble-averaged MSD, which considers all *N* particles, is given by:

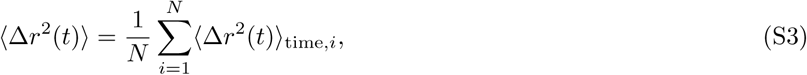

or equivalently:

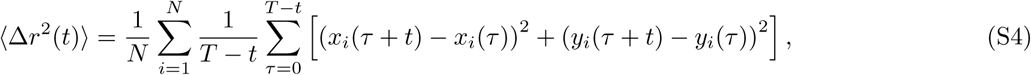

where *N* is the total number of particles. The exponent, *α*, was obtained by fitting the obtained time-averaged MSD to a power law for *t* = 0.005 s to *t* = 0.05 s for a given particle.

### Growth curve measurements

Growth curves were measured by turbidity measurements of growing cell cultures. Cell cultures were aliquoted into wells of a multiwell plate. We used a TECAN Spark Multimode Microplate Reader to measure the optical density with incubation and shaking during growth in the well plate.

### Modelling Trehalose Dependent Selection

We model the freeze-thaw adaptation by assuming trehalose produced at stationary phase to be the primary phenotypic marker. We assume that the growth rate of cells, the probability of a cell going into quiescence in stationary phase, the probability of survival under a freeze - thaw cycle, as well as the duration of lag time before re-entry into the cell cycle, depend on this stationary phase trehalose concentration.

We start with a population of *N* individual cells whose intracellular trehalose amount in stationary phase, given by *τ*, is drawn initially from a Gaussian distribution with mean *τ*_0_ and a narrow peak of with standard deviation 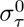. We take the simplest assumption for the dependence of each cell’s growth rate (or fecundity per unit time in the simulation) on stationary phase trehalose concentration,

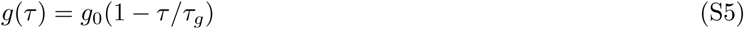

where *g*_0_ is the highest achievable growth rate and *τ_g_* the amount of trehalose that inhibits growth to effectively zero. At each time step in the growth process, we calculate the number of offspring for each cell according to the above equation, and add them to the population. Each offspring is also subject to a small mutation, which acts to change the trehalose concentration of the offspring with respect to its parent. This mutational effect on offspring trehalose is drawn from a Pearson Distribution at a mutation rate *m* for each offspring. The distribution has a mutational standard deviation *σ_m_*, a kurtosis *k_m_* and no skew. Thus, most mutations have no effect upon the offspring’s trehalose, preserving the value inherited from the parent cell. Simultaneously, some large effect mutations, both positive and negative are possible.

For each division, we assume a constant amount of a finite resource is consumed. This continues till the resource runs out and the population faces resource exhaustion. Upon starvation, yeast forms a heterogeneous population of quiescent and non-quiescent cells. We model the switch to quiescence as a stochastic process, where the probability of each cell going into quiescence is given by

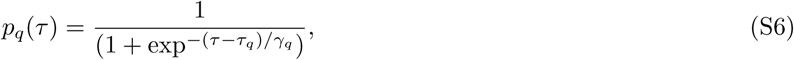

where *τ_q_*is the trehalose concentration at which probability of quiescence is 50% and *γ_q_* is a parameter quantifying how quickly the probability rises to 1 as *τ* increases. For each cell we decide the fate of the cell by comparing this probability with a randomly generated number.If the probability given by S6 is higher than the random number the cell goes into a quiescent state. Upon freezing, the non-quiescent population entirely perishes. Among the quiescent cells, the cells with trehalose concentration above a cut off survive, since trehalose has been shown to be correlated with higher survival to stress even in exponential phase [58]. We model the probability of survival,

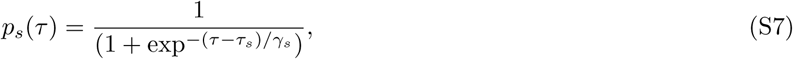

where *τ_s_* is the trehalose concentration at which probability of survival under freeze thaw is 50% and *γ_s_* is a parameter quantifying how quickly the probability rises to 1 as *τ* increases. In general we assume that *γ_s_ < γ_q_*and *τ_s_ > τ_g_*since the freezing process is a harsher stress than nutrient scarcity in stationary phase. Thus, both quiescence and the freeze thaw process shift the mean trehalose of the population to higher values since cells with lower trehalose amounts perish.

When the surviving population is allowed to thaw and re-introduced into a growth medium, the cells with higher trehalose concentrations have a shorter lag time, and begin to grow earlier as trehalose is crucial for re-entry into the cell cycle [39]. Longer lag times mean less resource available for growth because a small amount of resource is consumed even in the lag phase and the amount depends on the length of the lag phase. We again assume a linear relationship between the lag time and trehalose amounts,

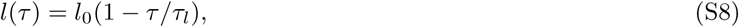

where *l*_0_ represents the maximum lag time possible, and *τ_l_* the trehalose value at which the lag time reduces to zero.

In each growth cycle, a higher trehalose value thus means a shorter lag phase, and more time available for growth, while simultaneously causing a lower growth rate. Lastly, we model the dilution process of the experimental setup by randomly selecting a fraction *d* of the population. This can introduce additional stochasticity into the evolution process.

At lower values of trehalose, the effect of a lower growth rate is not sufficient to stop the increase in mean trehalose of the population caused by a shorter lag phase. Cells with higher trehalose are able to produce a larger number of offspring and take over the population.

As the population undergoes another round of growth and glucose exhaustion, we once again have a heterogeneous population, with the distribution of quiescent cells now having a higher mean amount of trehalose production. A larger proportion of the cells are now able to survive the freeze-thaw process because of this upward shift. Hence, our simulation includes both a component of trehalose-dependent stochastic switching to quiescence, as well as growth and lag mediated selection for higher trehalose levels. The trade-off between the decrease in growth rate and the positive effect on all other factors from increasing trehalose sets the final equilibrium value of trehalose in the simulations. The effect of a lower growth rate begins to dominate at larger trehalose values, causing the mean trehalose of the population to stagnate.

Since the increase in the mean trehalose for the population is driven entirely by new beneficial mutations, the trajectory of individual replicates shows many jumps in the mean trehalose. The averaging over several replicates smoothens out the curve since the mutations, and the resulting sweep of the population can occur at different time points. Parameter values used in our simulation can be found in Table III.

**TABLE III.** Parameter values for simulations.

|  |  |
| --- | --- |
| N | $5 \times 10^4$ |
| $\tau_0$ | 500 |
| $\sigma_\tau^0$ | 1 |
| $\sigma_m$ | 2 |
| $k_m$ | 200 |
| $l_0$ | 6 |
| $\tau_l$ | 800 |
| $g_0$ | 10 |
| $\tau_g$ | 1200 |
| $\gamma_q$ | 100 |
| $\tau_q$ | 500 |
| $\gamma_s$ | 50 |
| $\tau_s$ | 700 |
| $d$ | 30 |

### Propagation of evolved populations without freeze-thaw stress

**FIG. S5.**
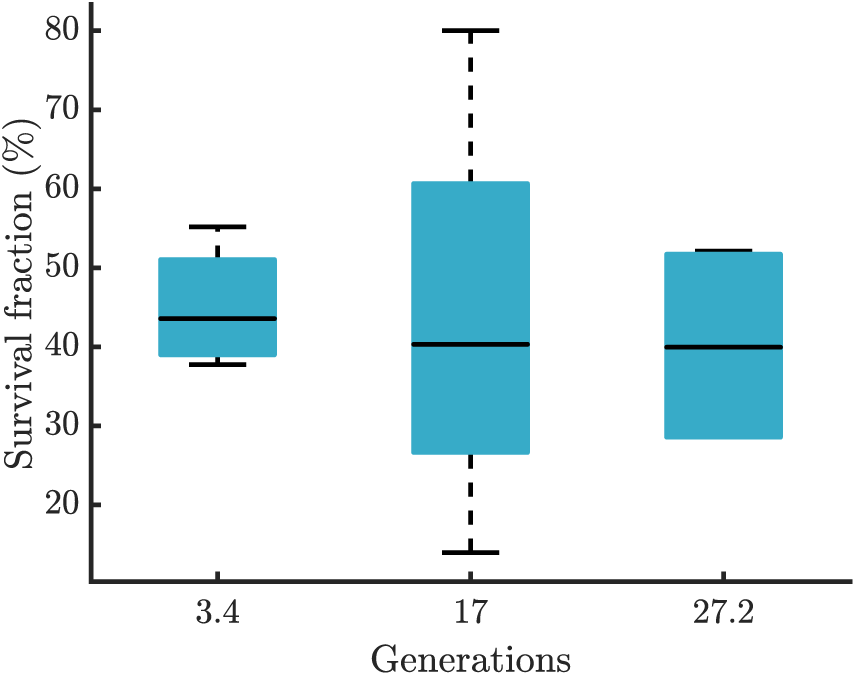
Average survival fraction of the one freeze-thaw evolved populations propagated without stress over several generations On each box, the central mark indicates the median, and the bottom and top edges of the box indicate the 25th and 75th percentiles, respectively. The whiskers extend to the most extreme data points not considered. outliers (there are no outliers)

All four replicates of the evolution experiment that selected for tolerance to one freeze-thaw were propagated without freeze-thaw stress for several generations to understand whether the tolerance could have been due to genetic inheritance. Populations were allowed to grow to stationary phase and then inoculated into fresh media and allowed to grow back to stationary phase again. This cycle was repeated several times, similar to the controls during the evolution. We found that the survival fraction does not change over more than 25 generations as shown in S5. This suggests genetic inheritance of freeze-thaw tolerance rather than epigenetic inheritance.

### Whole genome sequencing

For each evolution experiment, the starter culture, evolved strains, and control populations were independently sequenced. The evolved and control samples were collected from the middle and end of the evolution experiment, as described in figure S6. 13 samples were sequenced for each evolution experiment. Samples aliquoted and stored at −80^◦^C during the evolution experiment were inoculated and grown as a liquid culture to stationary phase. DNA from cells of these cultures was isolated using a Phenol:Chloroform:Isoamylalcohol extraction method. A DNA library was generated from this using the library preparation kit - Illumina® DNA Prep, (M) Tagmentation (Catalog no. - 20018705). The library was then amplified by PCR and sequenced on the NovaSeq 6000 platform using an SP flowcell with a sequencing read length of 2×100bp to obtain paired-end reads for all 26 samples.

**TABLE IV.** Sequenced samples

| Sample | Description |
| --- | --- |
| CTPO | Parent strain for one freeze-thaw tolerance selection |
| CTPT | Parent strain for three freeze-thaw tolerance selection |
| CTC1M1 | Control sample 1 of one freeze-thaw selection at mid-point of evolution |
| CTC1M2 | Control sample 2 of one freeze-thaw selection at mid-point of evolution |
| CTC3M1 | Control sample 1 of three freeze-thaw selection at mid-point of evolution |
| CTC3M2 | Control sample 2 of three freeze-thaw selection at mid-point of evolution |
| CTF1M1 | Replicate 1 of one freeze-thaw selection at mid-point of evolution |
| CTF1M2 | Replicate 2 of one freeze-thaw selection at mid-point of evolution |
| CTF1M3 | Replicate 3 of one freeze-thaw selection at mid-point of evolution |
| CTF1M4 | Replicate 4 of one freeze-thaw selection at mid-point of evolution |
| CTF3M1 | Replicate 1 of three freeze-thaw selection at mid-point of evolution |
| CTF3M2 | Replicate 2 of three freeze-thaw selection at mid-point of evolution |
| CTF3M3 | Replicate 3 of three freeze-thaw selection at mid-point of evolution |
| CTF3M4 | Replicate 4 of three freeze-thaw selection at mid-point of evolution |
| CTC1E1 | Control sample 1 of one freeze-thaw selection at end-point of evolution |
| CTC1E2 | Control sample 2 of one freeze-thaw selection at end-point of evolution |
| CTC3E1 | Control sample 1 of three freeze-thaw selection at end-point of evolution |
| CTC3E2 | Control sample 2 of three freeze-thaw selection at end-point of evolution |
| CTF1E1 | Replicate 1 of one freeze-thaw selection at end-point of evolution |
| CTF1E2 | Replicate 2 of one freeze-thaw selection at end-point of evolution |
| CTF1E3 | Replicate 3 of one freeze-thaw selection at end-point of evolution |
| CTF1E4 | Replicate 4 of one freeze-thaw selection at end-point of evolution |
| CTF3E1 | Replicate 1 of three freeze-thaw selection at end-point of evolution |
| CTF3E2 | Replicate 2 of three freeze-thaw selection at end-point of evolution |
| CTF3E3 | Replicate 3 of three freeze-thaw selection at end-point of evolution |
| CTF3E4 | Replicate 4 of three freeze-thaw selection at end-point of evolution |

### Sequencing analysis

#### s. Quality checks on sequencing output

The initial quality of sequencing reads was checked using FastQC. Three of the 11 tests—sequence content, GC content, and duplication level—returned quality failure or warning for all 52 sample files.

#### t. Preparation of reads for mapping

Trimmomatic was used to filter and trim reads prior to mapping. Bases were trimmed from either end if their quality was below 20. Additionally, the first 20 bases were removed regardless of their quality. Illuminaclip command was used to remove adapter sequence contaminants from read sequences. Trimmomatic first searches for a near exact match of a short adapter segment, of a maximum of 16 bp, within reads and then attempts to extend such “seed” sequences. A maximum of two mismatches were permitted in the seed sequences. A score of 30 (∼50 bases match) was required to consider the read a palindromic match (when the read-through was detectable on both reads of a pair) and a score of 10 for a direct match. The minimum adapter length required to be detected was set to 2 in the palindromic mode. Both reads were retained after the clipping. Reads of a minimum length of 60 after clipping were retained. The adapter sequence used was “CTGTCTCTTATACACATCT”, specific to the sequencing platform. The following command was run to apply the above-mentioned filters:

~~~
TrimmomaticPE -threads <NCPU> -phred33 -trimlog <LOG-file> <READ-1> <READ-2> -baseout <READ-out>
*‹→* ILLUMINACLIP:<ADAPTER-file>:2:30:10:2:True LEADING:20 TRAILING:20 HEADCROP:20 MINLEN:60
~~~

#### u. Mapping of reads to reference

The diploid strain CEN.PK113-7D a/*α*, Assembly accession: GCA 002571405.2 ASM257140 [73] was used as the reference genome to map the filtered and trimmed reads. The BWA-MEM algorithm was used to align paired-end reads to the reference genome with default parameters. Option -M was used to mark short split hits as secondary alignments that were to be ignored by downstream tools. Summary statistics for alignment and depth of coverage for every reference position were collected using samtools; flagstat and depth, respectively.

**FIG. S6.**
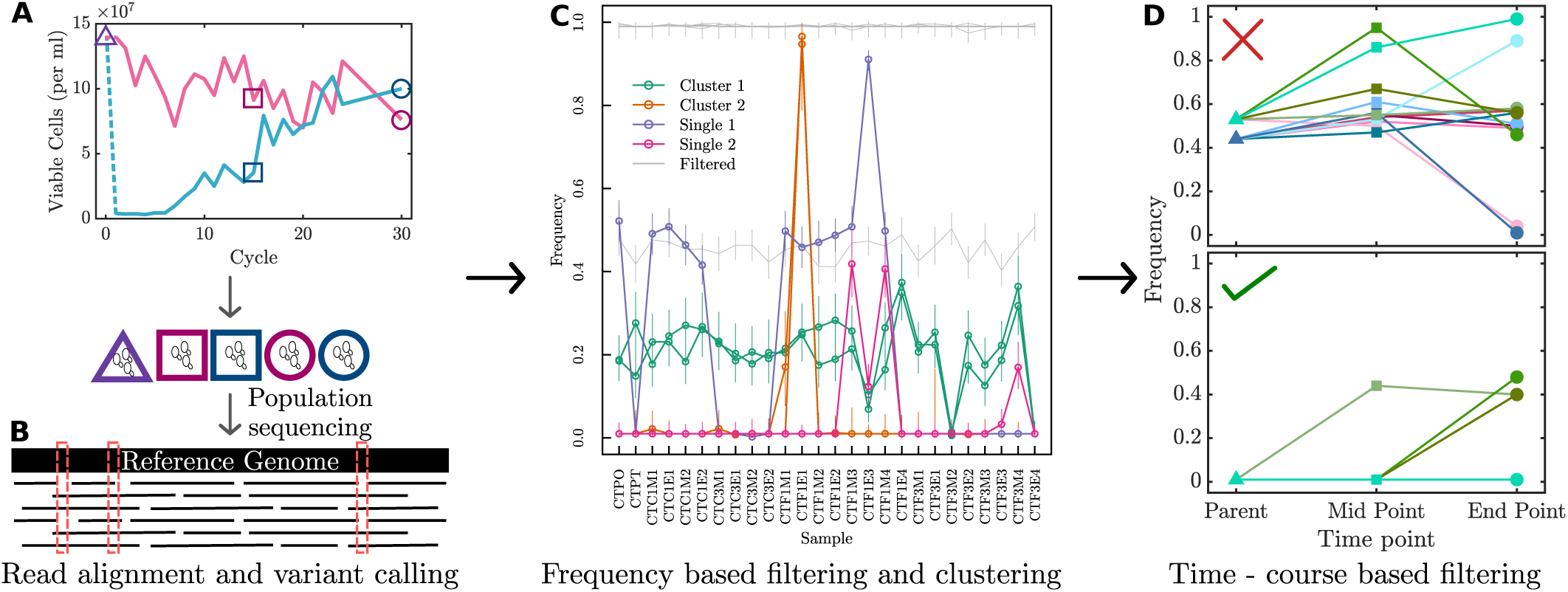
Sketch of the analysis of sequencing reads. **A** Representation of samples collected for sequencing. Sample collection points are displayed on a plot of the the number of viable cells in the control and evolving population across freeze-thaw-growth cycles. **B** Mapping of reads to a reference genome followed by variant calling. **C** A few representative variants filtered and clustered based on their frequency across all samples. Error bars denote 95% confidence intervals. **D** Trajectories of frequencies of representative variants from parent strain till the end point of selection. Each line represents a sample.

~~~
bwa mem -M -t <NCPU> -R “@RG\tID:ScerFTE\tSM:<SAMPLE>\tPL:ILLUMINA” <READ-1> <READ-2> |
*‹→* samtools sort -@ <NCPU> -o <sorted.bam> -
samtools flagstat --threads <NCPU> <sorted.bam> > <summary.flagstat>
samtools depth <sorted.bam> > <cov.txt>
~~~

#### v. Variant calling

SNPs and Indel variants were called for all samples together using bcftools mpileup with both minimum base quality and mapping quality set to 20, with option {per-sample-mF (minimum number and fraction of gapped reads) to increase the sensitivity of indel detection. Additionally, the-redo-BAQ option was used since BAQ (Base Alignment Qualities) has been proposed as a solution to find SNPs around indels, as an alternative to realignment around indels [74]. Base qualities (BQ) for variant calls were replaced with min{BQ, BAQ}. A maximum of 1000 reads per input alignment file was used for a position. The multi-allelic caller was used, with a ploidy of 2.

~~~
bcftools mpileup --redo-BAQ --min-BQ 20 --min-MQ 20 --per-sample-mF --annotate
*‹→* “FORMAT/AD,FORMAT/ADF,FORMAT/ADR,FORMAT/DP,FORMAT/SP,INFO/AD,INFO/ADF,INFO/ADR” --threads
*‹→* <NCPU> -d 1000 -f -b <sorted.bam> -Ou | bcftools call -mv --threads <NCPU> --ploidy 2
*‹→* -Ob -o <variants.bcf>
~~~

#### w. Variant filtering

All filtering operations were performed using bcftools. First, the SNP and Indel variants were separated. Appropriate thresholds for various variant quality statistics were derived by treating the variants shared by both parental samples as true variants and the variants private to either parent as false variants. This was done in the following steps:

Variants of either type present in parental samples were extracted using the following command:

~~~
bcftools view -s <sample1,sample2> -a --min-ac 1
~~~

Option -a was used to remove alternative alleles from ALT not seen in the subset. Option {min-ac was used to only print the line with at least 1 non-reference allele remaining. The following command listed variants present in only 1 sample *i.e.*, private variants:

~~~
bcftools view -s <SAMPLE1> -x -v <snps/indels>
~~~

Option -x was used to print sites where only the subset of samples have a non-ref allele. Option -v was used to extract SNP or Indel variant types.

Further, the private variant files for the parental samples CTC1P1 and CTC3P1 were merged. This list was subtracted from the collection of all parental variants to get a list of variants shared by both parents. Variants that were mutant homozygous in both samples were extracted and used as true variants (182 SNPs + 186 Indels). Heterozygous variants were extracted from the set of private variants and used as false variants (143 SNPs + 142 Indels).

~~~
bcftools merge <sample1.vcf.gz> <sample2.vcf.gz>
bcftools isec -p <SHARED> -c all -C <parental.vcf.gz> <private.vcf.gz>
bcftools view -i ’COUNT(GT=“AA”)=2’ <SHARED>/0000.vcf
bcftools view -i ’COUNT(GT=“RA”)=1’ <private.vcf>
~~~

The following thresholds were used based on variant quality statistics: SNPs:

- QUAL >= 50 (Variant Quality)
- DP <= 20000 (Depth)
- MQB > 1E-20 (Mapping Quality Bias)
- BQB > 1E-20 (Base Quality Bias)
- RPB > 1E-20 (Read Position Bias)
- SP <= 50 (Strand Bias)

Indels:

- QUAL >= 50 (Variant Quality)
- IDV >= 20 (Number of Supporting Reads)
- IMF >= 0.1 (Fraction of Supporting Reads)

The variants were filtered based on these thresholds using the following commands:

~~~
bcftools filter -i ’QUAL>=50 && INFO/DP<=20000 && RPB>1e-20 && MQB>1e-20 && BQB>1e-20’
*‹→* <snps_raw_all.vcf> | bcftools filter -e ’FMT/SP>50’
bcftools filter -i ’QUAL>=50 && IDV>=20 && IMF>=0.1’ <indels_raw_all.vcf>
~~~

Variants passing the above filters were then separated by experiments, and samples were rearranged based on time. Indels separated by only 3 or fewer bases were filtered out (filter -G 3).

#### x. Frequency-based filtering

Up to three non-ref alleles were present for Indels. Frequencies for each non-reference allele were extracted and analyzed separately. For SNPs, only one site with more than one non-reference allele was found for both experiments. The second non-reference allele at the SNP site in the 1-Freeze Thaw experiment was present only in one of the control samples. The SNP site with more than one non-reference allele in the 3-Freeze Thaw experiment had low depth of coverage, exceeding 10 in only one sample with only one or two reads for the alternate allele.

To obtain frequencies of the first non-ref allele for each variant, we used the following query:

~~~
bcftools query -f “%CHROM %POS %REF[ %AD{1}/%DP]\n” <snps_pass.vcf>
~~~

For others with n alleles:

~~~
bcftools view -i ’N_ALT>=n’ <indels_pass.vcf> | bcftools query -f “%CHROM %POS %REF[
*‹→* %AD{n}/%DP]\n”
~~~

A 95% confidence interval for each frequency estimate was obtained to determine whether a significant change occurred in the frequency of a variant in any sample during the course of the evolution experiment. Given the observed frequency of the variant in a sample of reads, the likelihood of different values for the true population frequency was evaluated assuming a binomial distribution of variant read counts. A minimum requirement of 10 total reads at a position was imposed to estimate frequency intervals. The minimum and maximum frequency values at which the probability of observed count falls below 5% constitute the limits of the confidence interval. Only the variants satisfying this condition in the reference as well as in the majority of samples were retained. Where the non-ref allele was already at almost 100% frequency in the parent, the frequency of the reference allele was followed instead. A mutant was removed if its frequency never exceeded 1%. A site was considered invariant if the upper estimate corresponding to its lowest frequency was greater than the lower estimate of its highest frequency. All invariant sites and sites where the minimum frequency change did not exceed 20% in any sample were excluded. The selected variant positions were pooled over experiments, the corresponding variants extracted from the raw VCF, and annotated using snpeff with a locally-built snpeff database for the reference assembly used in this study.

#### y. Frequency-based clustering of variants

Multiple variants may be present in the same cellular lineage, in which case their frequencies would be expected to change together across samples. Using the confidence intervals on frequencies as estimated above, a distance was calculated between pairs of sites as the sum of the minimum difference between estimated frequencies over samples. Pairs with distance ¡ 0.04 were considered as members of the same cluster. If a variant that occurred across clusters was associated with different variants across clusters, the associated variants were expected to be insignificantly contributing to the selected phenotype. Hence, independent variants or variants that were independently a part of one or more clusters were selected as the final variants.

#### z. Frequency-trajectory-based clustering of variants

The trajectories of the frequency of each variant were assessed across the parent, mid-point, and final samples. The variants for which the frequency trajectory fluctuated were filtered out. Only those variants whose frequency monotonically increased or decreased across the trajectory were retained.

*aa. Analysis of complementarity of genotypes across replicates* Using the Saccharomyces Genome Database (SGD), the final list of variants across all samples were checked for interactions with other variants. The possibility of one variant being a regulator of another was also checked using SGD.

*bb. Estimation of the population frequency of a gene-duplicated mutant* Copy number variation is another genotypic variant that could affect phenotype. Therefore, the genome was analysed for changes in copy numbers of genes. The coverage for each gene of each sample was used to estimate the change in copy number of the gene. The following model was used to estimate the frequency of copy number variants in a population, assuming that all copy number variants are gene duplications.

Regions of a genome are sequenced to varying depths due to sequencing bias. Let *x* be the per-base depth of coverage to the reference of a gene that’s present in a single copy in the sample. If an additional copy of the gene is present in the sample in the same region of the genome, then the expected coverage is 2*x*. However, if the second copy is in a different region which is represented in the sequencing data at a factor r of the mean coverage of the original region, then the total expected coverage of the duplicated gene is *x + rx*.

The above expression would be sufficient if the genome of a single clone with two copies of the gene was sequenced, or if the entire population carried the duplicated gene. Instead of single clones, we have sequenced samples of the whole population, and we have seen in the SNP analysis that some key mutants are present in the majority of the population but not in the entire population. Therefore, we must account for the possibility that the additional copy of the gene is only present in a fraction *f* of the population, which makes the expression for the expected coverage to be *x* (1+ *rf*), where *x* is the expected coverage of the gene in the absence of duplication.

The depth of sequencing coverage is expected to vary from sample to sample, leading to variation in the coverage of any target gene across samples even in the absence of duplication. Therefore, a normalization of gene coverage to sample coverage is required. Coverage of a gene normalized to median coverage of its chromosome for a sample should be similar across samples in the absence of gene duplication. In particular, we derived the expected coverage of a gene in a sample in the absence of duplication (*x* _ij_) by using the relation where *x* _i0_ is the observed coverage of gene *i* in the parent sample and *x* _m-_ is the median coverage of the chromosome of gene *i* in the parent or in sample *j*. With this, the expected coverage of a gene in a sample in case of a duplication event is *x* _ij_(1+*rf*).

The expected coverage of a duplicated gene, based on the above model, can be equated with its observed coverage in sample, from which, a measure of *rf* can be obtained as (Obs – Exp) / Exp. In this expression, the population frequency *f* of a gene-duplicated mutant is confounded with the variation in coverage across chromosomal regions, measured with *r*. Since a gene-specific estimate of *r* cannot be obtained based on coverage alone, we estimated the range of *r* from gene-to-gene variation in normalized depth of coverage seen in parent samples assuming that they carry no duplicated genes relative to the reference. The 95 % Confidence Interval of *r* based on parent samples was (0.7,1.5). Given the expression for *rf*, lower and upper estimates of population frequency of a gene-duplicated mutant in a sample can be obtained by using the upper and lower bounds on *r* respectively. The median of these two estimates is taken to be the central estimate of the population frequency *f*. Note that while *r* can exceed 1, *f* must be restricted to the closed interval [0,1]. Thus, estimates of population frequency less than 0 or greater than 1 were replaced with 0 or 1 respectively.

**TABLE V.**
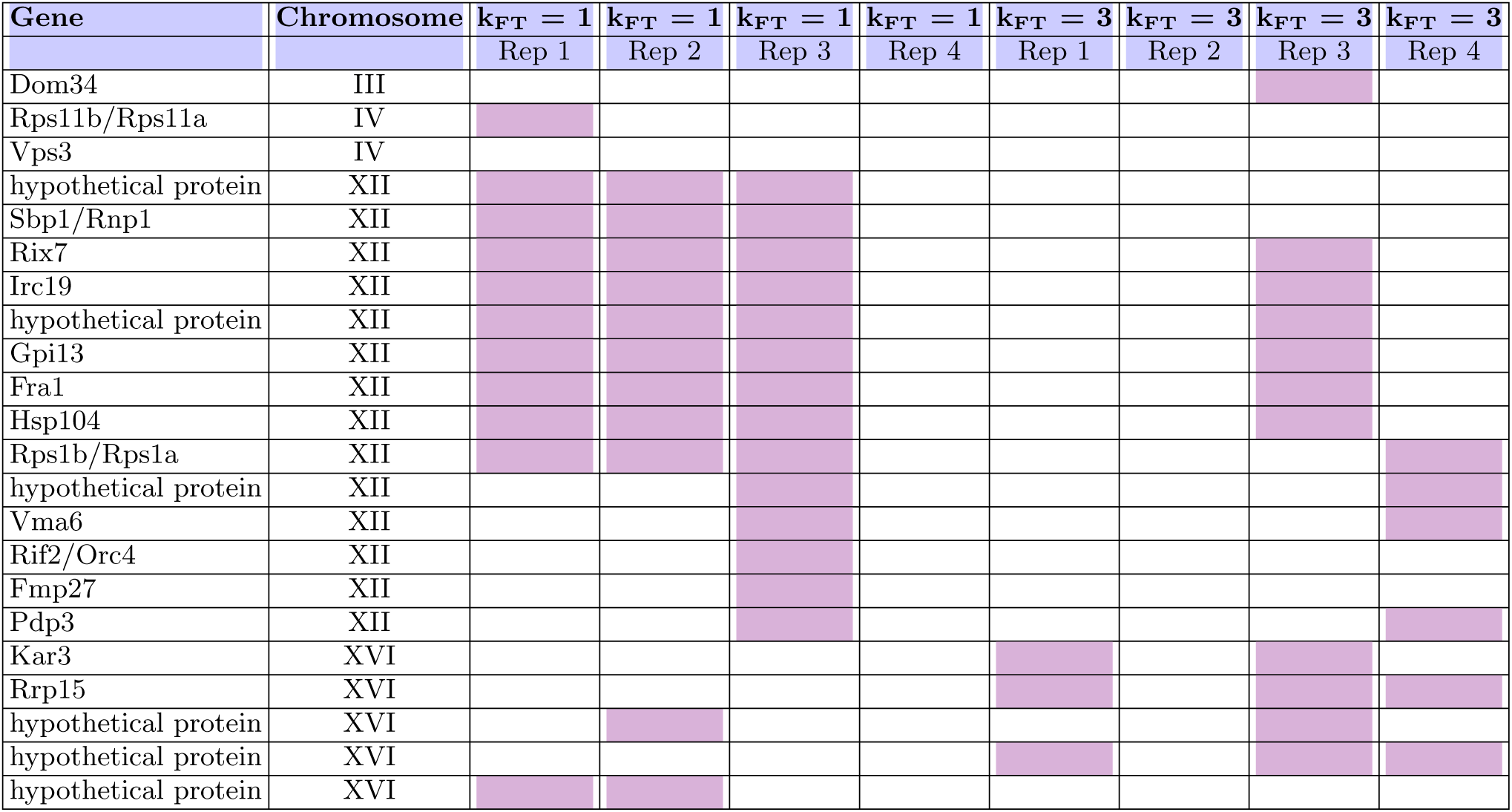
Replicates of populations of cells adapted to freeze-thaw stress do not have any duplicated genes in common. List of duplicated genes across all replicates including both the k_FT_ = 1 and k_FT_ = 3 selection regimes. Coloured cells denote a duplication.

## Criteria used for calling gene duplication events

1. At least 20 reads mapped to the gene in the parent sample.
2. At least 20 reads mapped in the majority of samples.
3. Lower estimate of population frequency (*f*) exceeds 0.5 in at least 1 sample.
4. The minimum change in frequency across samples, calculated as the difference between the highest lower estimate and the lowest upper estimate of frequency, is at least 0.2.

## References

[1] F. Franks. Biophysics and Biochemistry at Low Temperatures. Cambridge University Press, 1985.

[2] P. Mazur. Cryobiology: The freezing of biological systems. Science, 168:939–949, 1970.

[3] T. J. Anchordoguy, A. S. Rudolph, J. F. Carpenter, and J. H. Crowe. Modes of interaction of cryoprotectants with membrane phospholipids during freezing. Cryobiology, 24:324–331, 1987.

[4] J. Wolfe and G. Bryant. Freezing, drying, and/or vitrification of membrane-solute-water systems. Cryobiology, 39:103–129, 1999.

[5] B. Halliwell and J. M. C. Gutteridge. Free radicals in biology and medicine. Oxford University Press, 2015.

[6] K. B. Storey and J. M. Storey. Molecular Mechanisms of Metabolic Arrest: Life in Limbo. Springer, 2017.

[7] K. L. Koster and A. C. Leopold. Sugars and desiccation tolerance in seeds. Plant Physiology, 171:995–1006, 2016.

[8] R. A. Fisher. The Genetical Theory of Natural Selection. Oxford University Press, 1930.

[9] M. Kimura. Evolutionary rate at the molecular level. Nature, 217:624–626, 1968.

[10] H. A. Orr. The genetic theory of adaptation: A brief history. Nature Reviews Genetics, 6:119–127, 2005.

[11] S. A. Kelly, T. M. Panhuis, and A. M. Stoehr. Phenotypic plasticity: Molecular mechanisms and adaptive significance. Comprehensive Physiology, 2:1417–1439, 2012.

[12] C. K. Ghalambor, L. B. Martin, and H. A. Woods. Assessment and the regulation of adaptive phenotypic plasticity. Development, 151:dev203101, 2023.

[13] M. J. West-Eberhard. Developmental Plasticity and Evolution. Oxford University Press, 2003.

[14] H. Tapia and D. Koshland. Trehalose is a versatile and long-lived chaperone for desiccation tolerance. Molecular Cell, 55:601–610, 2014.

[15] Juan Carlos Argüelles. Physiological roles of trehalose in bacteria and yeasts: a comparative analysis. Archives of Microbiology, 174(4):217–224, 9 2000. URL: http://link.springer.com/10.1007/s002030000192 http://www.ncbi.nlm.nih.gov/pubmed/11081789, doi:10.1007/s002030000192.

[16] Claudio de Virgilio. The essence of yeast quiescence. FEMS Microbiology Reviews, 36(2):306–339, 2012. doi:10.1111/j.1574-6976.2011.00287.x.

[17] J. H. Crowe, L. M. Crowe, and D. Chapman. Preservation of membranes in anhydrobiotic organisms: The role of trehalose. Science, 223:701–703, 1984.

[18] E. Eleutherio, R. S. Araujo, A. G. Oliveira, and M. D. Pereira. The role of trehalose in yeast stress response. FEMS Yeast Research, 15:fov056, 2015.

[19] Matthias Christoph Munder, Daniel Midtvedt, Titus Franzmann, Elisabeth Nuske, Oliver Otto, Maik Herbig, Elke Ulbricht, Paul Müller, Anna Taubenberger, Shovamayee Maharana, Liliana Malinovska, Doris Richter, Jochen Guck, Vasily Zaburdaev, and Simon Alberti. A pH-driven transition of the cytoplasm from a fluid-to a solid-like state promotes entry into dormancy. eLife, 5:e09347, 3 2016. URL: https://elifesciences.org/articles/09347, doi:10.7554/eLife.09347.

[20] B. R. Parry, I. V. Surovtsev, M. T. Cabeen, and, et al. The bacterial cytoplasm has glass-like properties and is fluidized during stress. Cell, 156:183–194, 2014.

[21] Maja M. Klosinska, Christopher A. Crutchfield, Patrick H. Bradley, Joshua D. Rabinowitz, and James R. Broach. Yeast cells can access distinct quiescent states. Genes and Development, 25(4):336–349, 2 2011. URL: http://genesdev.cshlp.org/lookup/doi/10.1101/gad.2011311, doi:10.1101/gad.2011311.

[22] Damien Laporte, Laure Jimenez, Läetitia Gouleme, and Isabelle Sagot. Yeast quiescence exit swiftness is influenced by cell volume and chronological age. Microbial Cell, 5(2):104–111, 2018. doi:10.15698/mic2018.02.615.

[23] Samuel B Leslie, Eitan Israeli, Bruce Lighthart, J. H. Crowe, and Lois M Crowe. Trehalose and sucrose protect both membranes and proteins in intact bacteria during drying. Applied and Environmental Microbiology, 61(10):3592–3597, 10 1995. URL: http://aem.asm.org/ http://www.ncbi.nlm.nih.gov/pubmed/7486995, http://www.pubmedcentral.nih.gov/articlerender.fcgi?artid=PMC167656 https://journals.asm.org/doi/10.1128/aem.61.10.3592-3597.1995, doi:10.1128/aem.61.10.3592-3597.1995.

[24] S. Ibneeva and T. Grinenko. Dissecting dormancy and quiescence in hematopoietic stem cells. Frontiers in Hematology, 10:1401713, 2024.

[25] C. H. Lee, C. C. Yu, B. Y. Wang, and W. W. Chang. Regulatory role of quiescence in the biological function of cancer stem cells. Stem Cell Reviews and Reports, 16:435–447, 2020.

[26] P. Mazur. Freezing of living cells: Mechanisms and implications. American Journal of Physiology, 247:C125–C142, 1984.

[27] D. Simonin, L. Beney, and P. Gervais. Sequence of biochemical events involved in cell death during dehydration and rehydration of *Saccharomyces cerevisiae*. Applied Microbiology and Biotechnology, 96:1117–1130, 2012.

[28] Yuta Koganezawa, Miki Umetani, Moritoshi Sato, and Yuichi Wakamoto. History-dependent physiological adaptation to lethal genetic modification under antibiotic exposure. Elife, 11:e74486, 2022.

[29] Sean C Sleight, Christian Orlic, Dominique Schneider, and Richard E Lenski. Genetic basis of evolutionary adaptation by Escherichia coli to stressful cycles of freezing, thawing and growth. Genetics, 180(1):431–43, 9 2008. URL: http://www.ncbi.nlm.nih.gov/pubmed/18757947 http://www.pubmedcentral.nih.gov/articlerender.fcgi?artid=PMC2535694, doi:10.1534/genetics.108.091330.

[30] Peter Mazur and Janice J Schmidt. Interactions of cooling velocity, temperature, and warming velocity on the survival of frozen and thawed yeast. Cryobiology, 5(1):1–17, 7 1968. URL: https://linkinghub.elsevier.com/retrieve/pii/S0011224068801385 http://www.ncbi.nlm.nih.gov/pubmed/5760041, doi:10.1016/s0011-2240(68)80138-5.

[31] Toshihide Nakamura, Hiroshi Takagi, and Jun Shima. Effects of ice-seeding temperature and intracellular trehalose contents on survival of frozen Saccharomyces cerevisiae cells. Cryobiology, 58(2):170–174, 2009. doi:10.1016/j.cryobiol.2008.11.012.

[32] Jun Shima, Akihiro Hino, Chie Yamada-Iyo, Yasuo Suzuki, Ryouichi Nakajima, Hajime Watanabe, Katsumi Mori, and Hiroyuki Takano. Stress Tolerance in Doughs of Saccharomyces cerevisiae Trehalase Mutants Derived from Commercial Baker’s Yeast. Applied and Environmental Microbiology, 65(7):2841–2846, 7 1999. URL: https://journals.asm.org/doi/10.1128/AEM.65.7.2841-2846.1999, doi:10.1128/AEM.65.7.2841-2846.1999.

[33] Anqi Chen and Patrick A. Gibney. Intracellular trehalose accumulation via the Agt1 transporter promotes freeze–thaw tolerance in Saccharomyces cerevisiae. Journal of Applied Microbiology, 133(4):2390–2402, 10 2022. URL: https://academic.oup.com/jambio/article/133/4/2390/6989000, doi:10.1111/jam.15700.

[34] Cihan Erkut, Vamshidhar R Gade, Sunil Laxman, and Teymuras V Kurzchalia. The glyoxylate shunt is essential for desiccation tolerance in C. elegans and budding yeast. eLife, 5:1–24, 4 2016. URL: http://elifesciences.org/lookup/doi/10.7554/eLife.13614 https://elifesciences.org/articles/13614, doi:10.7554/eLife.13614.

[35] Ali Eroglu, Michael J Russo, Robert Bieganski, Alex Fowler, Stephen Cheley, Hagan Bayley, and Mehmet Toner. Intra-cellular trehalose improves the survival of cryopreserved mammalian cells. Nature Biotechnology, 18(2):163–167, 2 2000. URL: https://www.nature.com/articles/nbt0200_163 http://biotech.nature.com, doi:10.1038/72608.

[36] J. Kruuv, J.R. Lepock, and A.D. Keith. The effect of fluidity of membrane lipids on freeze-thaw survival of yeast. Cryobiology, 15(1):73–79, 2 1978. URL: https://linkinghub.elsevier.com/retrieve/pii/0011224078900093, doi:10.1016/0011-2240(78)90009-3.

[37] Yuhui Li, Hao Wang, and Pan Tingrui. Intracellular ice formation (IIF) during freeze-thaw repetitions. International Journal of Heat and Mass Transfer, 64:436–443, 2013. URL: 10.1016/j.ijheatmasstransfer.2013.04.036, doi:10.1016/j.ijheatmasstransfer.2013.04.036.

[38] P. H. Calcott and R. A. MacLeod. The survival of Escherichia coli from freeze–thaw damage: the relative importance of wall and membrane damage. Canadian Journal of Microbiology, 21(12):1960–1968, 12 1975. URL: http://www.nrcresearchpress.com/doi/10.1139/m75-284, doi:10.1139/m75-284.

[39] Lei Shi, Benjamin M. Sutter, Xinyue Ye, and Benjamin P. Tu. Trehalose Is a Key Determinant of the Quiescent Metabolic State That Fuels Cell Cycle Progression upon Return to Growth. Molecular Biology of the Cell, 21(12):1982–1990, 6 2010. URL: https://www.molbiolcell.org/doi/10.1091/mbc.e10-01-0056, doi:10.1091/mbc.e10-01-0056.

[40] Isabelle Sagot and Damien Laporte. The cell biology of quiescent yeast – a diversity of individual scenarios. Journal of Cell Science, 132(1), 1 2019. URL: https://journals.biologists.com/jcs/article/132/1/jcs213025/3037/The-cell-biology-of-quiescent-yeast-a-diversity-of, doi:10.1242/jcs.213025.

[41] Siyu Sun and David Gresham. Cellular quiescence in budding yeast. Yeast (Chichester, England), 38(1):12–29, 1 2021. URL: https://onlinelibrary.wiley.com/doi/10.1002/yea.3545 http://www.ncbi.nlm.nih.gov/pubmed/33350503 http://www.pubmedcentral.nih.gov/articlerender.fcgi?artid=PMC8208048, doi:10.1002/yea.3545.

[42] Oliver Otto, Philipp Rosendahl, Alexander Mietke, Stefan Golfier, Christoph Herold, Daniel Klaue, Salvatore Girardo, Stefano Pagliara, Andrew Ekpenyong, Angela Jacobi, Manja Wobus, Nicole Töpfner, Ulrich F. Keyser, Jörg Mansfeld, Elisabeth Fischer-Friedrich, and Jochen Guck. Real-time deformability cytometry: on-the-fly cell mechanical phenotyping. Nature methods, 12(3):199–202, 3 2015. URL: http://www.ncbi.nlm.nih.gov/pubmed/25643151, doi:10.1038/nmeth.3281.

[43] M. Delarue, G.P. Brittingham, S. Pfeffer, I.V. Surovtsev, S. Pinglay, K.J. Kennedy, M. Schaffer, J.I. Gutierrez, D. Sang, G. Poterewicz, J.K. Chung, J.M. Plitzko, J.T. Groves, C. Jacobs-Wagner, B.D. Engel, and L.J. Holt. mTORC1 Controls Phase Separation and the Biophysical Properties of the Cytoplasm by Tuning Crowding. Cell, 174(2):338–349, 7 2018. URL: https://linkinghub.elsevier.com/retrieve/pii/S0092867418306548, doi:10.1016/j.cell.2018.05.042.

[44] W. P. Voth. Yeast vectors for integration at the HO locus. Nucleic Acids Research, 29(12):59e–59, 2002. doi:10.1093/nar/29.12.e59.

[45] Viktor M. Boer, Christopher A. Crutchfield, Patrick H. Bradley, David Botstein, and Joshua D. Rabinowitz. Growth-limiting Intracellular Metabolites in Yeast Growing under Diverse Nutrient Limitations. Molecular Biology of the Cell, 21(1):198–211, 1 2010. URL: https://www.molbiolcell.org/doi/10.1091/mbc.e09-07-0597, doi:10.1091/mbc.e09-07-0597.

[46] Markus Basan, Tomoya Honda, Dimitris Christodoulou, Manuel Hörl, Yu-Fang Fang Chang, Emanuele Leoncini, Avik Mukherjee, Hiroyuki Okano, Brian R. Taylor, Josh M. Silverman, Carlos Sanchez, James R. Williamson, Johan Paulsson, Terence Hwa, and Uwe Sauer. A universal trade-off between growth and lag in fluctuating environments. Nature, 584(7821):470–474, 8 2020. URL: 10.1038/s41586-020-2505-4 https://www.nature.com/articles/s41586-020-2505-4, doi:10.1038/s41586-020-2505-4.

[47] Linda L. Breeden and Toshio Tsukiyama. Quiescence in Saccharomyces cerevisiae. Annual Review of Genetics, 56:253–278, 2022. doi:10.1146/annurev-genet-080320-023632.

[48] Xia Wang, Kotaro Fujimaki, Geoffrey C. Mitchell, Jungeun Sarah Kwon, Kimiko Della Croce, Chris Langsdorf, Hao Helen Zhang, and Guang Yao. Exit from quiescence displays a memory of cell growth and division. Nature Communications, 8(1):321, 8 2017. URL: 10.1038/s41467-017-00367-0, https://www.nature.com/articles/s41467-017-00367-0 http://www.ncbi.nlm.nih.gov/pubmed/28831039 http://www.pubmedcentral.nih.gov/articlerender.fcgi?artid=PMC5567331, doi:10.1038/s41467-017-00367-0.

[49] Anita Panek. Function of trehalose in Baker’s yeast (Saccharomyces cerevisiae). Archives of Biochemistry and Biophysics, 100(3):422–425, 3 1963. URL: https://linkinghub.elsevier.com/retrieve/pii/0003986163901073, doi: 10.1016/0003-9861(63)90107-3.

[50] Chris Allen, Sabrina Buttner, Anthony D. Aragon, Jason A. Thomas, Osorio Meirelles, Jason E. Jaetao, Don Benn, Stephanie W. Ruby, Marten Veenhuis, Frank Madeo, and Margaret Werner-Washburne. Isolation of quiescent and nonquiescent cells from yeast stationary-phase cultures. The Journal of Cell Biology, 174(1):89–100, 7 2006. URL: http://www.ncbi.nlm.nih.gov/pubmed/16818721 http://www.pubmedcentral.nih.gov/articlerender.fcgi?artid=PMC2064167 https://rupress.org/jcb/article/174/1/89/53700/Isolation-of-quiescent-and-nonquiescent-cells-from, doi: 10.1083/jcb.200604072.

[51] Monika Opalek, Bogna Smug, Michael Doebeli, and Dominika Wloch-Salamon. On the Ecological Significance of Phenotypic Heterogeneity in Microbial Populations Undergoing Starvation. Microbiology Spectrum, 10(1), 2 2022. URL: https://journals.asm.org/doi/10.1128/spectrum.00450-21, doi:10.1128/spectrum.00450-21.

[52] L. Diniz-Mendes, E. Bernardes, P. S. de Araujo, A. D. Panek, and V. M. F. Paschoalin. Preservation of frozen yeast cells by trehalose. Biotechnology and Bioengineering, 65(5):572–578, 12 1999. URL: https://onlinelibrary.wiley.com/doi/10.1002/(SICI)1097-0290(19991205)65:5%3C572::AID-BIT10%3E3.0.CO;2-7, doi:10.1002/(SICI)1097-0290(19991205)65:5<572::AID-BIT10>3.0.CO;2-7.

[53] Tomas Alarcon and Henrik Jeldtoft Jensen. Quiescence: A mechanism for escaping the effects of drug on cell populations. Journal of the Royal Society Interface, 8(54):99–106, 2011. doi:10.1098/rsif.2010.0130.

[54] Anthony D. Aragon, Angelina L. Rodriguez, Osorio Meirelles, Sushmita Roy, George S. Davidson, Phillip H. Tapia, Chris Allen, Ray Joe, Don Benn, and Margaret Werner-Washburne. Characterization of Differentiated Quiescent and Nonquiescent Cells in Yeast Stationary-Phase Cultures. Molecular Biology of the Cell, 19(3):1271–1280, 3 2008. URL: https://www.molbiolcell.org/doi/10.1091/mbc.e07-07-0666, doi:10.1091/mbc.e07-07-0666.

[55] Felix Keller, Maja Schellenberg, and Andres Wiemken. Localization of trehalase in vacuoles and of trehalose in the cytosol of yeast (Saccharomyces cerevisiae). Archives of Microbiology, 131(4):298–301, 6 1982. URL: http://link.springer.com/ 10.1007/BF00411175, doi:10.1007/BF00411175.

[56] T Chen, J P Acker, A Eroglu, S Cheley, H Bayley, A Fowler, and M Toner. Beneficial effect of intracellular trehalose on the membrane integrity of dried mammalian cells. Cryobiology, 43(2):168–81, 2001. URL: http://www.ncbi.nlm.nih.gov/pubmed/11846471, doi:10.1006/cryo.2001.2360.

[57] Laura B. Persson, Vardhaan S. Ambati, and Onn Brandman. Cellular Control of Viscosity Counters Changes in Temperature and Energy Availability. Cell, 183(6):1572–1585, 12 2020. URL: https://linkinghub.elsevier.com/retrieve/pii/S0092867420313805, doi:10.1016/j.cell.2020.10.017.

[58] Hugo Tapia, Lindsey Young, Douglas Fox, Carolyn R. Bertozzi, and Douglas Koshland. Increasing intracellular trehalose is sufficient to confer desiccation tolerance to Saccharomyces cerevisiae. Proceedings of the National Academy of Sciences, 112(19):6122–6127, 5 2015. URL: http://www.ncbi.nlm.nih.gov/pubmed/25918381, http://www.pnas.org/lookup/doi/10.1073/pnas.1506415112, http://www.pubmedcentral.nih.gov/articlerender.fcgi?artid=PMC4434740 https://pnas.org/doi/full/10.1073/pnas.1506415112, doi:10.1073/pnas.1506415112.

[59] Ryan P Joyner, Jeffrey H Tang, Jonne Helenius, Elisa Dultz, Christiane Brune, Liam J Holt, Sebastien Huet, Daniel J Müller, and Karsten Weis. A glucose-starvation response regulates the diffusion of macromolecules. eLife, 5:1–26, 3 2016. URL: https://elifesciences.org/articles/09376, doi:10.7554/eLife.09376.

[60] Ye Won Kwon, Jae-Han Bae, Seul-Ah Kim, and Nam Soo Han. Development of Freeze-Thaw Tolerant Lactobacillus rhamnosus GG by Adaptive Laboratory Evolution. Frontiers in Microbiology, 9(NOV):1–10, 11 2018. URL: https://www.frontiersin.org/article/10.3389/fmicb.2018.02781/full, doi:10.3389/fmicb.2018.02781.

[61] Peter Mazur. Physical and Temporal Factors Involved in the Death of Yeast at Subzero Temperatures. Biophysical Journal, 1(3):247–264, 1 1961. URL: https://linkinghub.elsevier.com/retrieve/pii/S0006349561868872, doi:10.1016/S0006-3495(61)86887-2.

[62] J. B. Griffiths, C. S. Cox, D. J. Beadle, C. J. Hunt, and D. S. Reid. Changes in cell size during the cooling, warming and post-thawing periods of the freeze-thaw cycle. Cryobiology, 16(2):141–151, 1979. doi:10.1016/0011-2240(79)90024-5.

[63] Tani Chen, Alex Fowler, and Mehmet Toner. Literature review: Supplemented phase diagram of the trehalose-water binary mixture. Cryobiology, 40(3):277–282, 5 2000. URL: https://linkinghub.elsevier.com/retrieve/pii/S0011224000922442, doi:10.1006/cryo.2000.2244.

[64] Siraje Arif Mahmud, Takashi Hirasawa, and Hiroshi Shimizu. Differential importance of trehalose accumulation in Saccharomyces cerevisiae in response to various environmental stresses. Journal of Bioscience and Bioengineering, 109(3):262–266, 2010. URL: 10.1016/j.jbiosc.2009.08.500, doi:10.1016/j.jbiosc.2009.08.500.

[65] Z. Petek Çakar, Urartu OS Seker, Candan Tamerler, Marco Sonderegger, and Uwe Sauer. Evolutionary engineering of multiple-stress resistant Saccharomyces cerevisiae. FEMS Yeast Research, 5(6-7):569–578, 4 2005. URL: https://academic.oup.com/femsyr/article-lookup/doi/10.1016/j.femsyr.2004.10.010, doi:10.1016/j.femsyr.2004.10.010.

[66] Nathalie Q Balaban, Jack Merrin, Remy Chait, Lukasz Kowalik, and Stanislas Leibler. Bacterial persistence as a phenotypic switch; Supplemental Materials. Science, 305(5690):1622–1625, 2004.

[67] O. Gefen, C. Gabay, M. Mumcuoglu, G. Engel, and N. Q. Balaban. Single-cell protein induction dynamics reveals a period of vulnerability to antibiotics in persister bacteria. Proceedings of the National Academy of Sciences, 105:6145–6149, 2008.

[68] K. J. Cheung and A. J. Ewald. Illuminating breast cancer invasion: Diverse roles for cell–cell interactions. Current Opinion in Cell Biology, 25:663–671, 2013.

[69] Monika Opalek, Hanna Tutaj, Adrian Pirog, Bogna J. Smug, Joanna Rutkowska, and Dominika Wloch-Salamon. A Systematic Review on Quiescent State Research Approaches in S. cerevisiae. Cells, 12(12):1608, 6 2023. URL: https://www.mdpi.com/2073-4409/12/12/1608, doi:10.3390/cells12121608.

[70] Dominika M. Wloch-Salamon, Katarzyna Tomala, Dimitra Aggeli, and Barbara Dunn. Adaptive Roles of SSY1 and SIR3 During Cycles of Growth and Starvation in Saccharomyces cerevisiae Populations Enriched for Quiescent or Nonquiescent Cells. G3 Genes—Genomes—Genetics, 7(6):1899–1911, 6 2017. URL: https://academic.oup.com/g3journal/article/7/6/1899/6029837, doi:10.1534/g3.117.041749.

[71] O. J. Rando and K. J. Verstrepen. Timescales of genetic and epigenetic inheritance. Cell, 128:655–668, 2012.

[72] Steven B. Haase and Steven I. Reed. Improved flow cytometric analysis of the budding yeast cell cycle. Cell cycle (Georgetown, Tex.), 1(2):117–121, 2002. doi:10.4161/cc.1.2.114.

[73] Alex N. Salazar, Arthur R. Gorter de Vries, Marcel van den Broek, Melanie Wijsman, Pilar de la Torre Cortes, Anja Brickwedde, Nick Brouwers, Jean Marc G. Daran, and Thomas Abeel. Nanopore sequencing enables near-complete de novo assembly of Saccharomyces cerevisiae reference strain CEN.PK113–7D. FEMS Yeast Research, 17(7):1–11, 2017. doi:10.1093/femsyr/fox074.

[74] Heng Li. Improving SNP discovery by base alignment quality. Bioinformatics (Oxford, England), 27(8):1157–8, 4 2011. URL: https://academic.oup.com/bioinformatics/article/27/8/1157/227268, http://www.ncbi.nlm.nih.gov/pubmed/21320865, http://www.pubmedcentral.nih.gov/articlerender.fcgi?artid=PMC3072548, doi:10.1093/bioinformatics/btr076.

